# Mitochondrial membrane proteins and VPS35 orchestrate selective removal of mtDNA

**DOI:** 10.1101/2021.08.06.455397

**Authors:** David Pla-Martin, Ayesha Sen, Sebastian Kallabis, Julian Nüchel, Kanjanamas Maliphol, Julia Hofmann, Marcus Krüger, Rudolf J. Wiesner

**Affiliations:** Center for Physiology and Pathophysiology, Institute of Vegetative Physiology, University of Köln, Köln, Germany; Cologne Excellence Cluster on Cellular Stress Responses in Aging-associated Diseases (CECAD), University of Köln, Köln, Germany; Center for Biochemistry, Faculty of Medicine, University of Köln, Köln, Germany; Center for Molecular Medicine Cologne, University of Köln, Köln, Germany

## Abstract

Integrity of mitochondrial DNA (mtDNA), encoding several subunits of the respiratory chain, is essential to maintain mitochondrial fitness. Mitochondria, as a central hub for metabolism, are affected in a wide variety of human diseases but also during normal ageing, where mtDNA integrity is compromised. Mitochondrial quality control mechanisms work at different levels, and mitophagy and its variants are critical to remove dysfunctional mitochondria together with mtDNA to maintain cellular homeostasis. Understanding the mechanisms governing a selective turnover of mutation-bearing mtDNA without affecting the entire mitochondrial pool is fundamental to design therapeutic strategies against mtDNA diseases and ageing. Here we show that mtDNA depletion after expressing a dominant negative version of the mitochondrial helicase Twinkle, or by chemical means, is due to an exacerbated mtDNA turnover. Targeting of nucleoids is controlled by Twinkle which, together with the mitochondrial transmembrane proteins ATAD3 and SAMM50, orchestrate mitochondrial membrane remodeling to form extrusions. mtDNA removal depends on autophagy and requires the vesicular trafficking protein VPS35 which binds to Twinkle-enriched mitochondrial subcompartments upon mtDNA damage. Stimulation of autophagy by rapamycin selectively removes mtDNA deletions which accumulated during muscle regeneration *in vivo*, but without affecting mtDNA copy number. With these results we unveil a new complex mechanism specifically targeting and removing mutant mtDNA which occurs outside the mitochondrial network. We reveal the molecular targets involved in a process with multiple potential benefits against human mtDNA related diseases, either inherited, acquired or due to normal ageing.

## INTRODUCTION

The accumulation over time of mutations in the mitochondrial genome (mtDNA) is a common process which has been shown to occur in many tissues ^1^ and is probably one of the hallmarks of aging ^2^. MtDNA is present in thousands of copies per cell, hence, impairment of mitochondrial function is observed only when the percentage of mutated mtDNA molecules surpasses a specific threshold^3^.

Cells possess a plethora of quality control mechanisms to survey the intactness of DNA, RNA and proteins, but also of entire organelles. In addition to bulk autophagy, which is responsible for the continuous and non-selective turnover of cellular material activated during nutrient shortage, specific mechanisms are initiated to remove malfunctioning organelles upon damage. The process of mitophagy has been investigated extensively in recent years as an important salvage pathway to remove dysfunctional mitochondria ^4^. The ongoing fission and fusion events of the dynamic mitochondrial network are important processes in order to survey mitochondrial quality by predisposing those parts of the network with impaired function to degradation. Mitochondrial fission is a requirement to initiate mitophagy of damaged mitochondria ^4^, whereas knockout of key players of mitochondrial fusion has been shown to induce mtDNA instability, either by causing a rapid accumulation of mtDNA alterations over time ^5^ or by mtDNA depletion ^6^.

Recently, a process with a high level of specificity involving mitochondrial derived vesicles (MDVs) was shown to remove not the complete organelle, but rather mitochondrial fragments containing specific cargo^7,8^. This mechanism requires the coordination of mitochondrial dynamics, mitophagy and also the vacuolar protein sorting (VPS) or retromer complex. In this process, changes in the mitochondrial membrane potential and the oxidation state of mitochondrial subcompartments induce the curvature of the membrane which is followed by recruitment of PINK1 and PARKIN ^9^. The retromer complex, formed by VPS26, VPS29 and VPS35, provides the force to generate a vesicle which is then delivered to lysosomes or peroxisomes in a process which is independent of the autophagy proteins ATG5 or LC3 ^10-12^. In summary, mitophagy and its variants are crucial pathways to degrade parts of the mitochondrial network, thus maintaining cellular fitness.

Many inherited forms of neurodegenerative diseases are examples for insufficient mitochondrial quality control. Mutations of specific receptors involved in targeting dysfunctional organelles to mitophagy like PINK1 and PARKIN, but also malfunction of lysosomal proteins ATP13A2 and LAMP3 among many others, as well as mutations in the retromer component *VPS35* ^13^ cause familial forms of Parkinson’
ss disease. Parkinson’s disease is caused by the specific degeneration of dopaminergic neurons which have been comprehensively shown to be a hotspot for the accumulation of large scale mtDNA deletions during normal aging ^14,15^ making mitochondrial quality control mechanisms especially important in these cells. Unfortunately, these mechanisms are not sufficient to counteract the often progressive phenotype of patients suffering from PD and other diseases caused by either maternally inherited or by acquired mtDNA alterations, the latter due to mutations in proteins essential for mtDNA replication and maintenance ^16^. Thus, therapeutic approaches to increase mitochondrial quality control and counteract the progression of mtDNA related diseases have been attempted ^17,18^, but a lack of specificity and activation of undesirable effects are still a hallmark ^19^. Therefore, understanding the specific mechanisms governing mtDNA turnover is pivotal to develop new therapeutic strategies against these syndromes.

Expressing the mitochondrial helicase Twinkle bearing dominant negative mutations causing mitochondrial disease in patients induces mtDNA instability ^20^. Expression of the disease related *PEO1* Twinkle mutation K320E (from now on K320E) in mouse models accelerates the accumulation of mtDNA deletions in postmitotic tissues like heart and muscle^21,22^ during aging and induce mtDNA depletion in proliferating cells such as epidermis and cartilage^23,24^. Recently, it has been shown that muscles from those mice accumulated deletions, but the main alterations were gene duplications, leading to a vast variety of different enlarged and rearranged molecules ^25^. Using a combination of *in vivo* and *in vitro* approaches, we have now identified the proteins involved in a new mechanism for targeting and specific degradation of mtDNA to avoid the accumulation of such mutations. Expression of K320E induces the formation of mitochondrial extrusions, much larger than MDVs, which are consecutively engulfed by autophagosomes and fused with lysosomes. Elimination of altered mtDNA molecules is preceded by the relocation of nucleoids to the poles of mitochondria, a process controlled by the interaction between the mitochondrial inner membrane protein ATAD3 and the nucleoid protein Twinkle. The translocase protein SAMM50 and VPS35 are essential to provide the required selectivity and specificity for mtDNA elimination. We show that stimulation of autophagy by rapamycin *in vivo* is sufficient to specifically remove deleted mtDNA, but without affecting the total mtDNA copy number. Thus, modulation of autophagy *in vivo* can be used as an approach to counteract the accumulation of mutations in mtDNA observed in several mitochondrial pathologies and during aging.

## RESULTS

### Accumulation of mtDNA mutations in skeletal muscles *in vivo* does not induce mitophagy

In a previous work we have found that, in extraocular muscles, which are first affected in *Peo1* patients, mtDNA alterations preferentially accumulated in fast twitch in contrast to slow oxidative fibers, indicating important differences in mitochondrial quality control mechanisms in different muscle fiber types ^22^. To study the nature of these mechanisms surveillant mtDNA integrity, we first analysed fast-twitch M. tibialis anterior (TA) and slow-oxidative M. soleus (SOL), muscles both rich in fibers with a high mitochondrial content in mice, but with a preferentially glycolytic (TA) vs. oxidative metabolism (SOL), respectively. As shown before by deep sequencing^25^, both muscles showed an accumulation of many mtDNA alterations in 24 months old animals (Fig. 1a). K320E mutant mice carry a wide variety of reorganized mtDNA molecules^25^, causing an inefficient amplification reaction leading to a smear of products, however there were no changes in total mtDNA copy number (Fig. 1b). Noteworthy, TA from aged wt mice also showed many mtDNA alterations, while only few were found in SOL. By conventional PCR, we analysed the presence of deletions common in aged mice ^22^ and selected a deletion covering about 4000bp (*Mus musculus* mtDNA-Δ^983-4977^), which was present in both mutant muscles (Fig. S1a). Considering that mtDNA copy number is on average 20% higher in SOL than in TA (Fig. S1b), we performed qPCR quantification using the D-Loop region as a reference and found that indeed, this deletion was on average 20 times more abundant in TA compared to SOL (Fig. 1c). Steady state levels of common mitochondrial autophagy markers showed that in SOL of K320E^skm^ mice, the general autophagy adaptors p62 and LC3 were significantly decreased while levels were similar in TA (Fig. 1d, e). Levels of the specific mitophagy adaptor Optineurin (OPT) and the mitochondrial marker TOM20 were similar in all samples. *In situ* immunofluorescence of LC3 and p62 confirmed reduced puncta of those proteins (Fig. S1c-f). Since such a reduction could reflect both an increase or a decrease of autophagy flux, we blocked autophagy flux by chloroquine, and analysed the conversion of LC3-I to LC3-II (Fig. 1f, g). We did not observe an increased autophagy flux in K320E mice compared to control mice. However, in SOL for K320E mice, chloroquine was not inducing a change in the LC3 ratio suggesting that, in this muscle, autophagy flux was already at maximum level in steady state.

**Figure 1.**
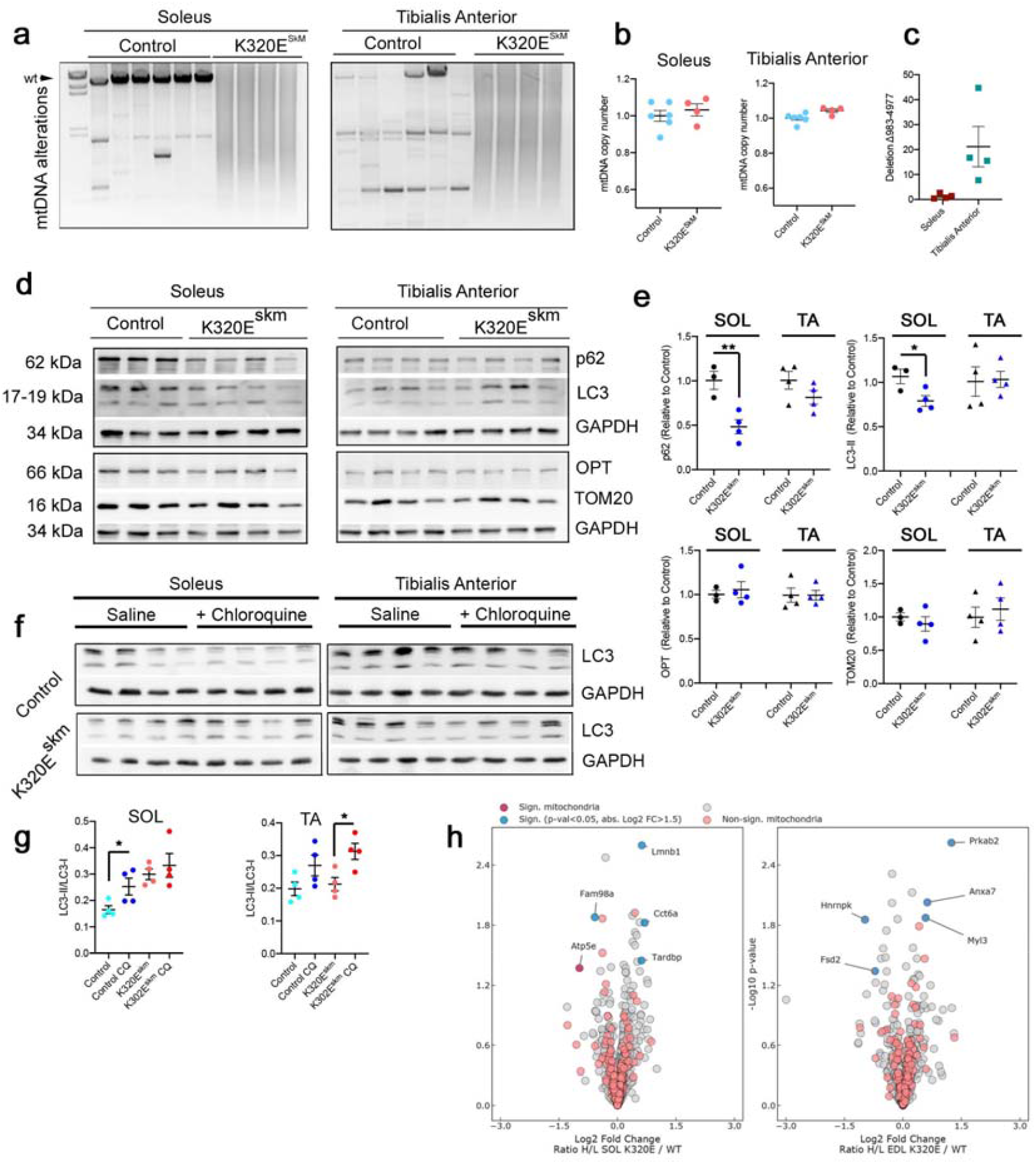
*In vivo* expression of Twinkle-K320E induces differential accumulation of mtDNA alterations in muscles. (**a**) Long range PCR analysis, (**b**) quantification of mtDNA copy number and, (**c**) qPCR quantification of deletion mtDNA-Δ983-4977 in M. soleus and M. Tibialis anterior from 24 months old control and K320E^SkM^ mice. (**d, e**) Western blot analysis and quantification of the indicated proteins in muscle extracts from 24 months old mice. M. soleus, control: n=3; Twinkle-K320E: n=4. M. tibialis anterior, control: n=4; Twinkle-K320E: n=4. (**f, g**) Western blot analysis and quantification of LC3 flux (LC3-II/LC3-I ratio) in muscle extracts from mice treated with saline or 50mg/kg chloroquine 4 hours before euthanasia. n=4 mice per treatment and genotype. Unpaired Students’
s T-test. Mean ± SEM. (**h**) Volcano plots for mitochondrial and non-mitochondrial proteins detected in the muscles analysed for *in vivo* Pulse SILAC. Red dots: mitochondrial proteins; dark red: mitochondrial proteins with significant difference; blue dots: non mitochondrial proteins with significance difference. (p-value < 0.05 and a H/L ratio fold change > 1.5). n=5.

We therefore hypothesized that oxidative fibres, which are most dependent on intact mitochondria, possess a faster mitochondrial turnover, hence maintaining mutated mtDNA molecules more efficiently below a pathogenic threshold. To visualize protein synthesis *in vivo*, we fed our mice with heavy lysine (^13^C_6_-lysine) for two weeks in order to get an estimate for mitochondrial turnover rates ^26^. Here, we selected SOL and M. extensor digitorum longus (EDL) for analysis, since the latter contains almost exclusively glycolytic fibers. We first confirmed that heavy Lysine incorporation was equal in all animals (Fig. S2a, b). We analysed H/L (heavy/light) ratios of detected proteins (1286 and 1224 for SOL, 1042 and 1018 for EDL in wild-type and K320E^skm^ mice, respectively) and filtered the mitochondrial proteins using Mitocarta 3.0 (Supplementary Table 1; Fig. S1c). Surprisingly, no mitochondrial protein was showing differential incorporation of heavy lysine comparing K320E mutant and wild-type animals, neither in EDL nor in SOL, suggesting that bulk mitochondrial turnover, here measured as an increase of mitochondrial biogenesis, is not enhanced in muscles bearing mtDNA alterations (Fig. 1h).

### Autophagy is required for depletion of mtDNA following oxidative damage

To further dissect the molecular pathways activated upon mtDNA instability, we generated stable C2C12 myoblast cell lines expressing tagged versions of Twinkle and K320E. Colocalization of these variants with mitochondria (outer membrane marker TOM20, Fig. S3a, b) as well as colocalization with mtDNA and the mtDNA binding protein TFAM was confirmed (Fig. S3c, d), showing that Twinkle is enriched in mitochondrial nucleoids.

To investigate if K320E expression leads to activation of autophagy or mitophagy in these cells, C2C12 myoblasts were additionally transfected with plasmids encoding LC3-GFP-mCherry or Fis1p-GFP-mCherry. Expression of K320E induced the accumulation of autolysosomes marked by LC3-GFP-mCherry (Fig. 2a, b; magenta signal). In agreement with our *in vivo* pSILAC results, expression of Fis1p-GFP-mCherry, a mitophagy reporter, showed no activation of acute mitophagy (Fig. 2c, d). Previous studies have demonstrated that the *in vitro* expression of several Twinkle missense mutations often leads to accumulation of mtDNA replication intermediates, producing mtDNA depletion ^27^. Consistently, expression of K320E leads to mtDNA depletion (Fig. 2e), but this was not related to a decrease in mtDNA replication rate, as observed by analysis of mtDNA replication foci in BrdU labelled cells (Fig. S4a, b). We hypothesized that accumulation of lysosomes and mtDNA depletion after expressing K320E was reflecting a novel selective mtDNA degradation process. Thus, we blocked lysosomal activity using chloroquine and showed that, indeed, under these conditions, mtDNA levels remained at control levels (Fig. 2f). Finally, we analysed the spatial relationship between Twinkle foci and autophagy structures, using LC3 as an autophagosome marker and Lamp1 as a late endosomal marker, respectively. As expected, most wt Twinkle foci distributed in a pattern reflecting the mitochondrial network and were independent of autophagy markers (Fig. 2g-j). However, colocalization with LC3 and LAMP1, respectively, was observed with K320E. The increased mtDNA degradation flux in K320E cells suggests that expression of this missense mutation harms mtDNA by unknown means. To visualize *in situ* mtDNA damage, we searched for the presence of 8-hydroxy-2’
sdeoxyguanosine (8-OHdG), a specific base modification induced by reactive oxygen species ^28^. We could not observe any specific staining with α-8-OHdG in steady state levels (Fig. S4c, d), however when cells were incubated in presence of chloroquine, we detected a specific accumulation of 8-OHdG decorating the mitochondrial network only in K320E expressing cells (Fig. 2k, l). In summary, our data indicate that K320E induces mtDNA depletion in a lysosome-dependent manner without involving acute mitophagy, pointing to a more specialized path.

**Figure 2.**
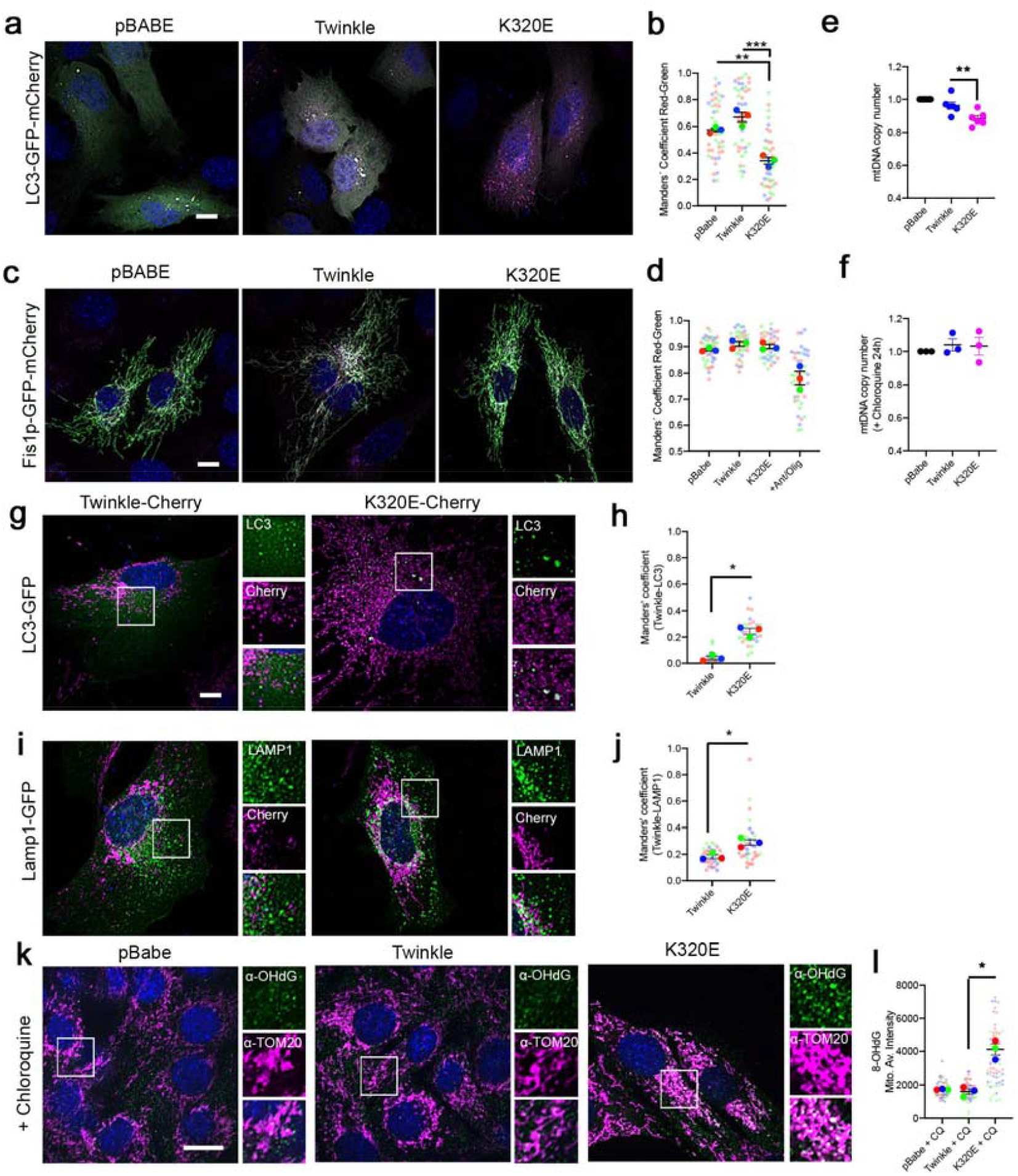
Twinkle-K320E triggers mtDNA damage and induces autolysosome accumulation independent of bulk mitophagy. (**a-e**) C2C12 expressing untagged Twinkle constructs transiently expressing the autophagy reporter LC3-GFP-mCherry or (**c**) the mitophagy reporter Fis1p-GFP-mCherry. Red signal shows lysosomal localization. (**b, c**) Manders’
s coefficient Red / Green quantification of transfected cells. n=3. A decrease in Manders’ coefficient indicates autolysosome or mito-lysosome accumulation, n=3, 10-15 cells per replicate. Cells treated overnight with 10µM Antimycin/Oligomycin were used to induce canonical mitophagy. (**e, f**) mtDNA copy number in C2C12 cells stably expressing Twinkle or Twinkle-K320E (K320E), respectively, vs. empty vector (pBABE). In (**f**), mtDNA levels were recovered by treating cells with the lysosomal inhibitor chloroquine for 24h, n=3. (**g-h**) Confocal images and quantification of C2C12 expressing mCherry tagged Twinkle transfected with plasmids encoding the autophagosome marker LC3-GFP and (**i, j**) lysosome marker Lamp1-GFP. Manders’ colocalization coefficient was used to confirm Twinkle colocalization with autophagic organelles, n=3 (>10 cells per experiment). (**k**) mtDNA oxidative damage detected by immunofluorescent labelling with α-OHdG and α-TOM20 in cells expressing Twinkle and treated with chloroquine (CQ) for 24h. (**l**) Relative intensity quantification of α-OHdG signal inside the mitochondrial network. Scale Bar, 10µm. (**c-f** and **l**) ANOVA, Tukey multiple comparison. **, p<0.01; ***, p<0.001. (**h** and **j**) Unpaired Student’s T-test. *, p<0.05. Mean ± SEM.

To further analyse the contribution of autophagy to mtDNA turnover, we turned to Atg5 KO MEFs and expressed Twinkle variants (Fig. 3a), since Atg5 has been shown to be important in quality control after mitochondrial damage ^29^. Analysis of mitochondrial nucleoids showed that K320E expression decreased nucleoid foci number in Atg5 wt cells (Fig. 3b). In contrast, in Atg5 KO cells, foci number was not altered by K320E expression (Fig. 3c). Nevertheless, expression of K320E reduced mtDNA copy number in both Atg5 WT and Atg5 KO cells (Fig. 3d, e). Analysis of mtDNA replication by BrdU labelling showed that, in contrast to wt cells, expression of K320E in autophagy deficient cells led to a reduced number of foci replicating mtDNA within the mitochondrial network (Fig. 3f, g). Chloroquine treatment also restored mtDNA copy number in Atg5 WT cells expressing K320E but not in Atg5 KO cells (Fig. 3h). Our data suggest that in contrast to wt cells where K320E induces mtDNA depletion linked to an increased nucleoid turnover rate, in autophagy deficient cells, mtDNA depletion is caused by reduced mtDNA replication, presumably to avoid accumulation of excessive mtDNA damage.

**Figure 3.**
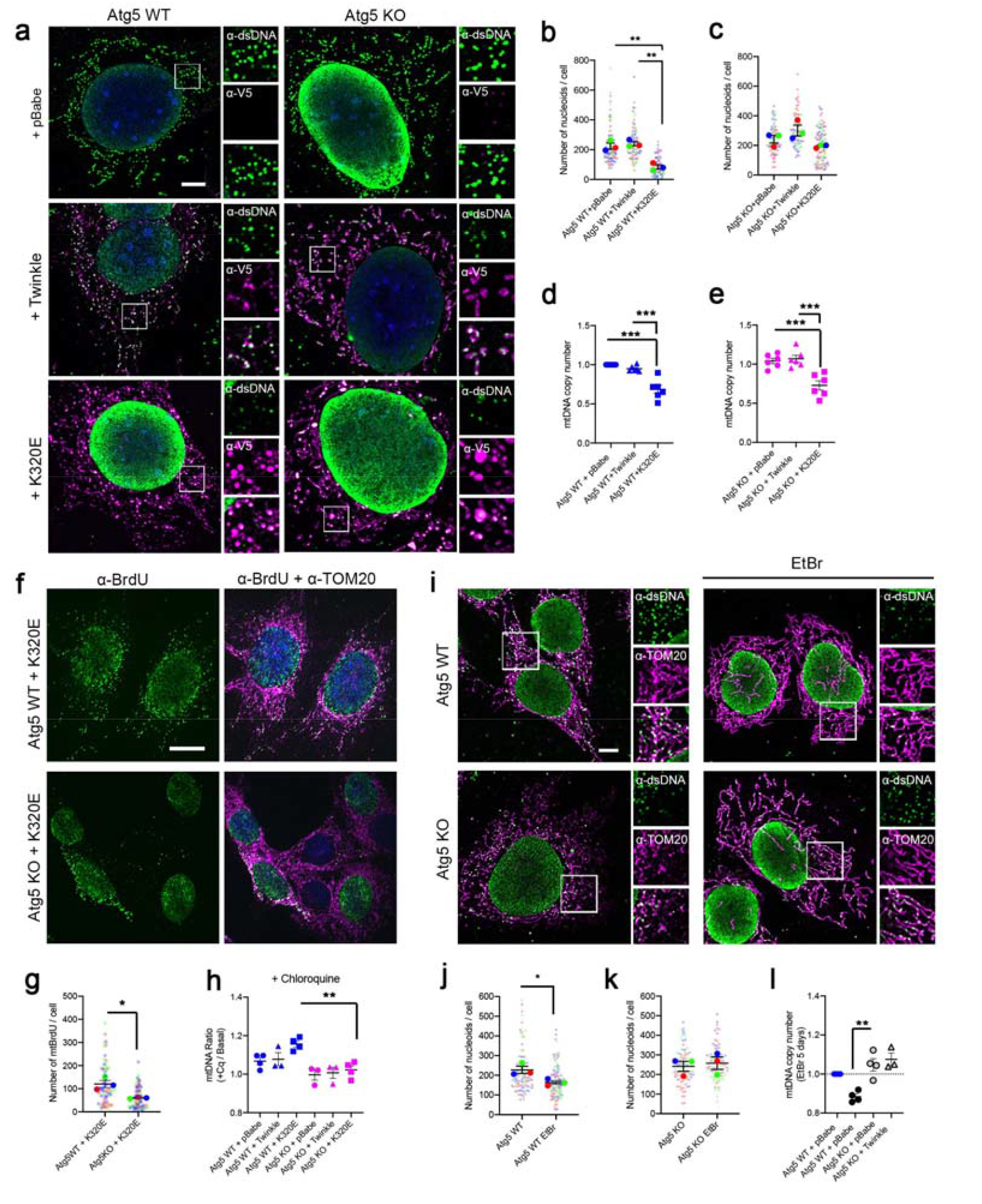
Autophagy is required for mtDNA clearance after mtDNA damage. Atg5 WT and KO cells were transduced with Twinkle-APEX-V5 plasmids. **(a)** α-DNA and α-V5-tag immunofluorescence confirming localization of Twinkle in mtDNA containing nucleoids. Scale bar 5µm. (**b, c**) Quantification of mtDNA foci number in Atg5 WT and Atg5 KO cells. n=3 (>25 cells per experiment). (**d, e**) Steady state mtDNA copy number in Atg5 cells. (**f, g**) α-BrdU and α-TOM20 immunofluorescence and quantification of mtDNA replicating foci detected by treating the cells for 6h with 20µM BrdU. n=3, >20 cells per replicate (**h**) Quantification of mtDNA copy number ratio in Atg5 cells treated with Chloroquine for 24h. n=3. (**i-k**) α-DNA and α-TOM20 immunofluorescence and mtDNA foci quantification in steady state and in cells treated with 50ng/ul EtBr for 7 days. n=3, >20 cells per replicate. (**l**) mtDNA copy number analysis for steady state and in cells treated for 7 days with EtBr. n=3-4. Scale Bar, 10µm. (**b-e, h** and **l**) ANOVA, Tukey multiple comparison. **, p<0.01; ***, p<0.001. (**g, j** and **k**) Unpaired Student’
ss T-test. *, p<0.05. Mean ± SEM.

Therefore, to evaluate the role of autophagy in mtDNA depletion, we induced mtDNA alterations by growing these cells for 7 days in medium containing Ethidium Bromide (EtBr), which we also found to provoke the accumulation of 8-OHdG within the mitochondrial network (Fig. S4c, e). Morphological analysis of mitochondrial nucleoids showed that EtBr decreases foci number in Atg5 WT cells but not in Atg5 KO (Fig. 3i-k). In line with this, Atg5 control cells showed consistent mtDNA depletion, whereas in Atg5 KO cells mtDNA copy number remained unchanged (Fig. 3l). This EtBr-induced depletion could not be recovered by overexpression of wt Twinkle, indicating again a prominent role of autophagy in mtDNA depletion in this case.

All these data together confirm that mtDNA damage induces activation of a mtDNA turnover mechanism, which is dependent on autophagosome formation and specifically degrades mtDNA but it does not activate an acute mitophagy response.

### mtDNA instability induces the formation of mitochondrial protrusions prior to lysosomal degradation

One of the most surprising features we noticed while studying cellular localization of Twinkle was that TOM20, a mitochondrial outer membrane marker, does not consistently overlap with Twinkle, a mitochondrial matrix protein (Fig. 4a and S3a, b). Live imaging microscopy using C2C12 clones expressing Twinkle-mCherry showed that occasionally, Twinkle moved to Mitotracker Green-negative poles of the network releasing a small particle which was also Mitotracker-negative (Fig. 4b, arrows). These particles were more evident in K320E expressing cells, where most of K320E was residing in such Mitotracker-negative structures (Fig. 4c). As Mitotracker green is independent of the inner membrane potential, we ruled out the possibility that these regions were depolarized areas of the mitochondrial network but mitochondrial regions with different cardiolipin composition.

**Figure 4.**
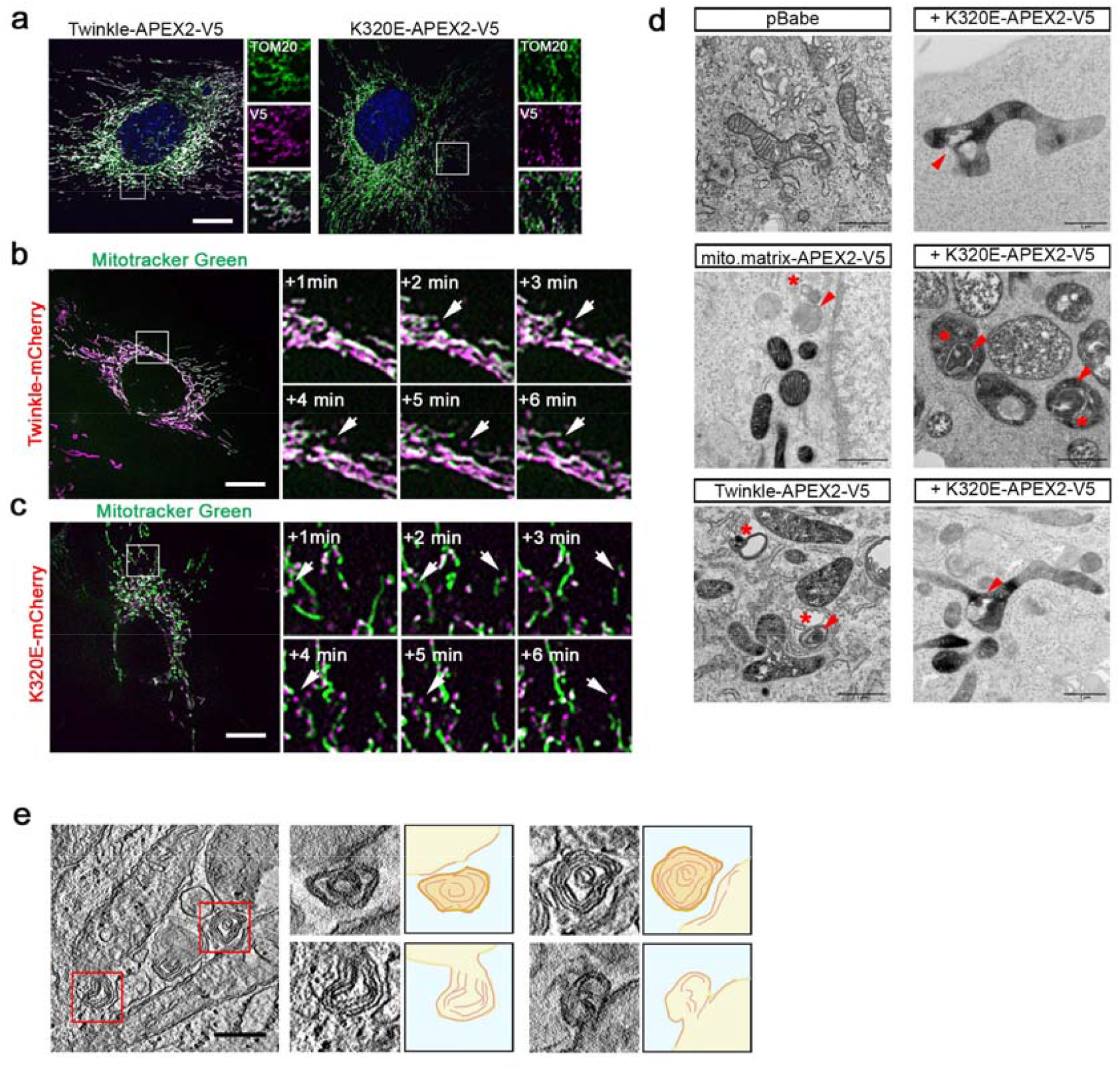
Twinkle-K320E localizes to specialized mitochondrial regions. (**a**) Immunofluorescence of C2C12 cells expressing Twinkle-APEX-V5 variants probed with α-TOM20 and α-V5 antibodies. Arrow shows TOM20 negative particle containing Twinkle-K320E. Bar, 10µm (**b, c**) Live imaging of C2C12 cells expressing tagged Twinkle-m-Cherry variants counterstained with mitotracker green. Insets represent a 1 min time lapse series of the selected area. Bar, 10µm (**d**) Transmission electron microscopic pictures of C2C12 cells expressing APEX2-V5 clones. APEX2 generates a black precipitate in presence of DAB. Red arrows indicate multimembrane structures, asterisks indicate autophagosomes. Bar, 1µm. (**e**) Electron tomography in C2C12 cells expressing Twinkle-K320E illustrating extrusion of mitochondrial fragments containing multimembrane structures. Bar, 0,5µm

To determine Twinkle localisation with high resolution, we used Twinkle-APEX2 fusion versions in C2C12 cell lines. APEX2 generates a black precipitate in the presence of DAB, which is visible by electron microscopy. Twinkle-APEX2 distribution in the matrix was heterogeneous and the organization of mitochondria and other cellular organelles was undisturbed by its expression (Fig. 4d). K320E-APEX2, however, accumulated in poles of the network, sometimes close to abnormal cristae structures (Fig. 3d, red arrows) and also inside lysosomes (Fig. 4d, red stars). Such multi-membrane structures were also observed in cells expressing a mitochondrial matrix targeted APEX2, but in that case, they did not show DAB precipitate (Fig. 4d, red stars). A very recent study demonstrated that mitochondrial fission occurring at mitochondrial poles is followed by mitophagy of mitochondrial compartments containing non-replicative mtDNA, and that this is preceded by mitochondrial cristae reorganization ^30^. Indeed, electron tomography of K320E expressing cells revealed that these structures are derived from the inner mitochondrial membrane and that they are formed by protrusions containing reorganized mitochondrial cristae (Fig. 4e).

### Twinkle interacts with membrane proteins to facilitate mtDNA removal

We speculated that mtDNA targeting prior to degradation is controlled by specific protein-protein interactions. Thus, as Twinkle follows mtDNA positioning and degradation through lysosomes, we sought to investigate how nucleoids containing Twinkle distribute in the mitochondrial matrix by analysing the Twinkle interactome. We performed immunoprecipitation of wt Twinkle fused to a V5-APEX2 epitope followed by mass spectrometry analysis. As expected, the majority of proteins interacting with Twinkle were related to mtDNA replication, transcription and translation (Supplementary Table 2), but only a few interactions were found to be significant (Fig. 5a, blue dots), one of them being ATAD3, an AAA-ATPase previously linked to anchoring and distribution of mtDNA to the inner membrane ^31^. Direct co-immunoprecipitation and immunofluorescence confirmed the physical interaction between ATAD3 and Twinkle (Fig. 5b; Fig. S5a) enlightening Twinkle as an important part of the link between mitochondrial nucleoids and the inner membrane.

**Figure 5.**
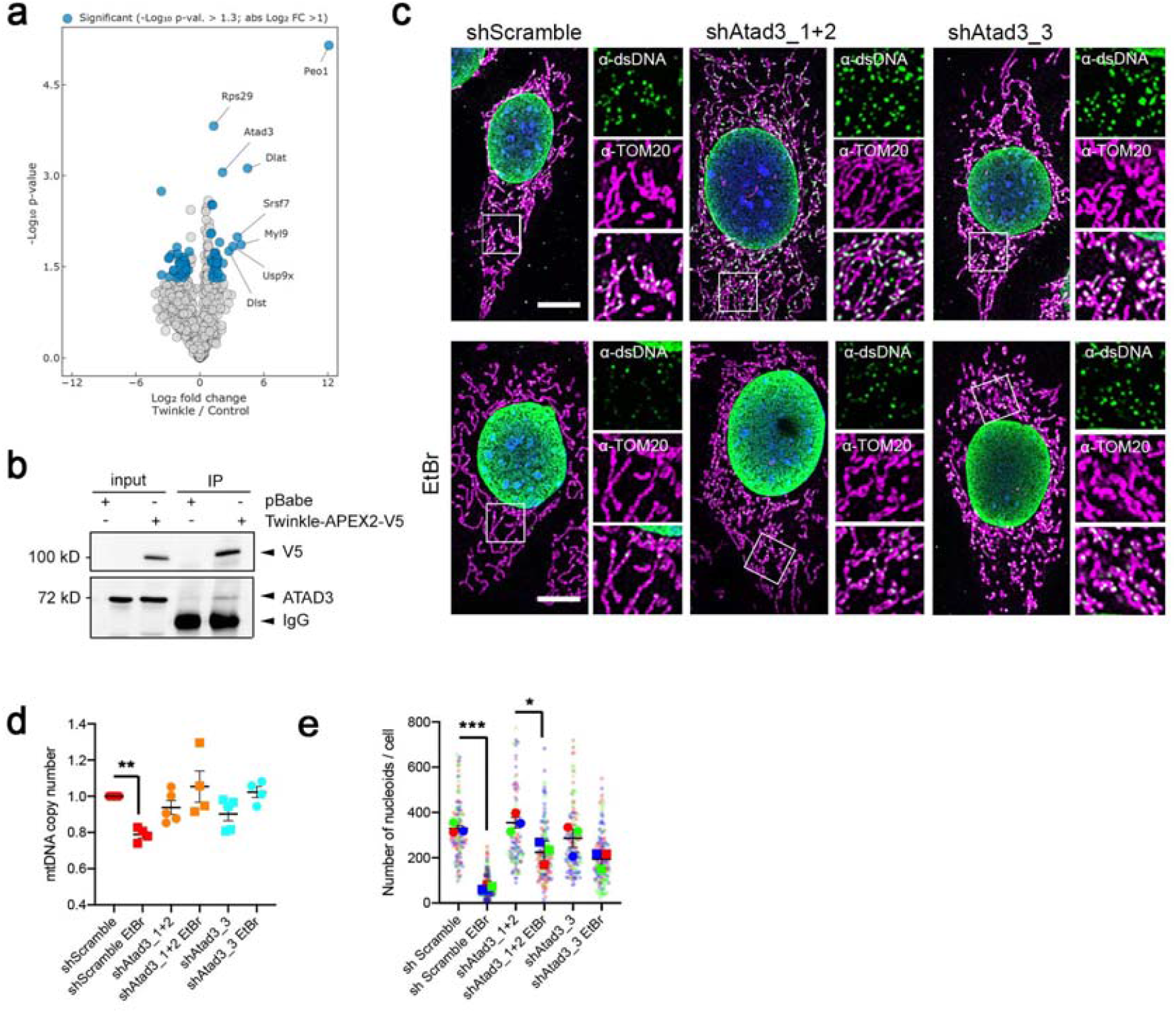
Twinkle mediates interaction of nucleoids with the inner mitochondrial membrane protein ATAD3. (**a**) Volcano plot showing proteins enriched after immunoprecipitation of Twinkle-APEX2-V5 in C2C12 cells. Differentially enriched proteins compared with cells transfected with empty vector (p-value < -0.05 and FC > 2 or < -2) are highlighted in blue (Peo1 = Twinkle). (**b**) Co-immunoprecipitation of V5-tagged Twinkle and ATAD3. (**c**) α-TOM20 and α-dsDNA immunofluorescence in Atad3 KD MEFs in steady state and grown for 7 days in presence of 50ng/ml EtBr. Bar, 10 µm. (**d**) Quantification of mtDNA copy number in steady state and EtBr treated cells. n=4-5. and (**e**) mtDNA foci quantification for Atad3 KD in steady state and EtBr treated cells. n=3, >20 cells per replicate. ANOVA, Tukey multiple comparison. *, p<0.05, **, p<0.01. Mean ± SEM.

To evaluate the role of these proteins in mtDNA turnover, we generated constitutive *Atad3* knock-down (KD) clones (shAtad3_1+2, shAtad3_3; Fig. S5b, c). *Atad3* KD clones showed steady state levels of mtDNA copy number comparable to control lines (Fig. 5c, d) and no changes in mitochondrial morphology (Fig. S5d). After one week of mild treatment with EtBr, control cells showed a persistent decrease of mtDNA copy number. However, this depletion was not observed in Atad3 KD clones (Fig. 5d). Number of nucleoid foci per cell was also higher in downregulated clones than in control cells upon EtBr treatment (Fig. 5e). These data suggest that nucleoid binding to the mitochondrial inner membrane through a Twinkle-ATAD3 interaction is essential for mtDNA elimination upon mtDNA damage.

### mtDNA turnover requires coordination of the retromer complex in a process independent of MDVs

We initially hypothesized that selective removal of mitochondrial fragments containing mtDNA was carried out through MDVs ^7^. The E3 Ubiquitin ligase MAPL has been shown to direct mitochondrial cargo to peroxisomes ^32^, while the endosomal adaptor Tollip specifically divert MDVs to lysosomes ^8^. In both cases, the VPS35-retromer complex initiates the force to generate a vesicle ^10^. Thus, in order to investigate if the specific mtDNA degradation we observed follows the MDV or a new pathway, we studied VPS35 and MAPL localization in Twinkle-mCherry expressing cells. Consistent with previous studies, expression of MAPL-GFP induced mitochondrial fission and formation of structures excluding TOM20 (Fig. S6a) ^32^. However, Twinkle was also excluded from those particles. Chloroquine treatment for 4h was sufficient to induce a rapid accumulation of Twinkle positive structures excluding TOM20 and MAPL, suggesting that the fate of those particles is the lysosomal compartment (Fig. S5b). In addition, we also corroborate that Twinkle colocalizes with VPS35 but excludes MAPL (Fig. S6c, d).

Next, we treated these cells as well with EtBr to induce mtDNA damage. One week of treatment was sufficient to reduce mtDNA copy number (Fig. S7a). Image quantification revealed that the percentage of VPS35 containing Twinkle was strongly increased upon EtBr treatment (Fig. 6a-c), even when the overall steady state levels of VPS35 structures did not change (Fig. 6d). Importantly, VPS35 also colocalized with dsDNA (Fig. 6e).

**Figure 6.**
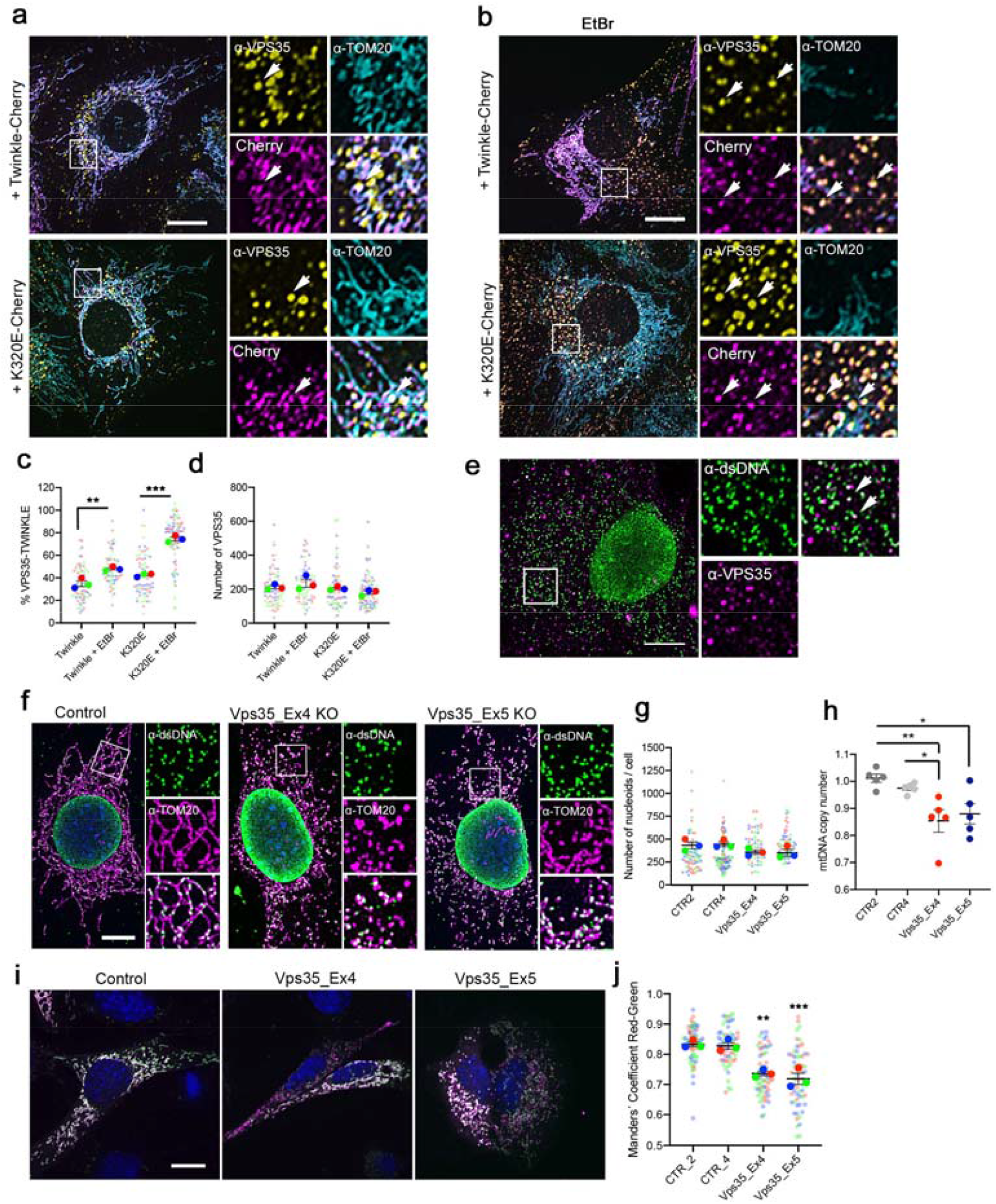
VPS35 is required for autophagy-dependent mtDNA removal. Immunofluorescence of C2C12 cells expressing Twinkle-mCherry variants labelled with α-VPS35 and α -TOM20 and grown in (**a**) basal medium or (**b**) treated for 7 days with 50ng/ml EtBr. Arrows indicate colocalization. Bar, 10µm. (**c, d**) Quantification of VPS35 particles in contact with Twinkle. n=3 (**e**) Immunofluorescence of C2C12 wild type cells labelled with α-dsDNA and α-VPS35. Arrows indicate colocalization between VPS35 and dsDNA puncta. Bar, 10µm. (**f**) α-TOM20 and α-dsDNA immunofluorescence of *Vps35* KO MEFs. Bar, 10µm. (**g**) mtDNA foci analysis (n=3) and (**h**) mtDNA copy number quantification of *Vps35* KO cells. n=5. (**i**) Control and *Vps35* KO cells transfected with Fis1p-GFP-mCherry plasmid to detect mitophagy. Red signal represents mito-lysosomes. Bar, 10µm. (**j**) Manders’
s coefficient quantification of transfected cells. n=3. A decrease in Manders’ coefficient indicates process activation. n=3 (20 images per replicate). ANOVA, Tukey multiple comparison. *, p<0.05; **, p<0.01; ***, p<0.001. Mean ± SEM

We then generated Vps35 KO clones in MEFs and selected two monoclonal lines targeting exon 4 (*Vps35*_Ex4) and exon 5 (*Vps35*_Ex5) (Fig. S6b, c). Vps35 KO cells showed no changes in mtDNA foci number but reduced levels of mtDNA copy number in steady state (Fig. 6f-h) and mitochondrial fragmentation (Fig. S7d). After 7 days of EtBr treatment, the number of mtDNA foci was further decreased (Fig. S7e, f) and mtDNA depletion was enhanced (Fig. S7g, h). We analysed the activation of canonical mitophagy in steady state by expressing Fis1p-GFP-mCherry (Fig. 6i) and found that, indeed, in Vps35 KO cells, acute mitophagy was activated (Fig. 6j). Moreover, VPS35 protein level was shown to be affected by lysosomal function, as chloroquine induced VPS35 accumulation in K320E cells (Fig. S7i, j) without interfering with late endosomes or mitochondrial content (Fig. S7i-n).

We hypothesize that VPS35 is necessary to fine-tune the elimination of mutated mtDNA without activating acute mitophagy and sought to investigate how VPS35 is recruited to mitochondria upon mtDNA damage. Thus, we performed VPS35 IP followed by MS analysis under basal conditions and in cells treated 7 days with EtBr (Fig. S8a). As expected, in basal medium the protein profile of VPS35 IP revealed poor interaction with mitochondrial proteins (Fig. 7a, red dots). Upon EtBr treatment, however, the association of mitochondrial proteins was markedly increased (Fig. 7b, c). Among them, we noticed the presence of mitochondrial membrane proteins such as VDAC1 and VDAC3, TIMMDC1 or TOMM40 as well as SAMM50, which interaction was confirmed by coimmunoprecipitation (Fig. 7d). The comparison of IP protein profiles of cells in basal vs. EtBr medium highlighted also a strong enrichment of the mAAA protease SPG7 (Fig. S8b). Remarkably, comparison between the IP protein profiles from Twinkle and K320E revealed also a strong enrichment of SAMM50 upon K320E immunoprecipitation (Fig. 7e). SAMM50 belongs to the sorting and assembly machinery in the mitochondrial intermembrane space and has important functions in the biogenesis of respiratory complexes, cristae morphology and mitochondrial shape ^33^. It was also described as a regulator of PINK1-Parkin mediated mitophagy but excluding the mtDNA for degradation ^34^. Immunoprecipitation of Twinkle confirms that, after mtDNA damage with EtBr, Twinkle can physically interact with SAMM50 (Fig. 7f).

**Figure 7.**
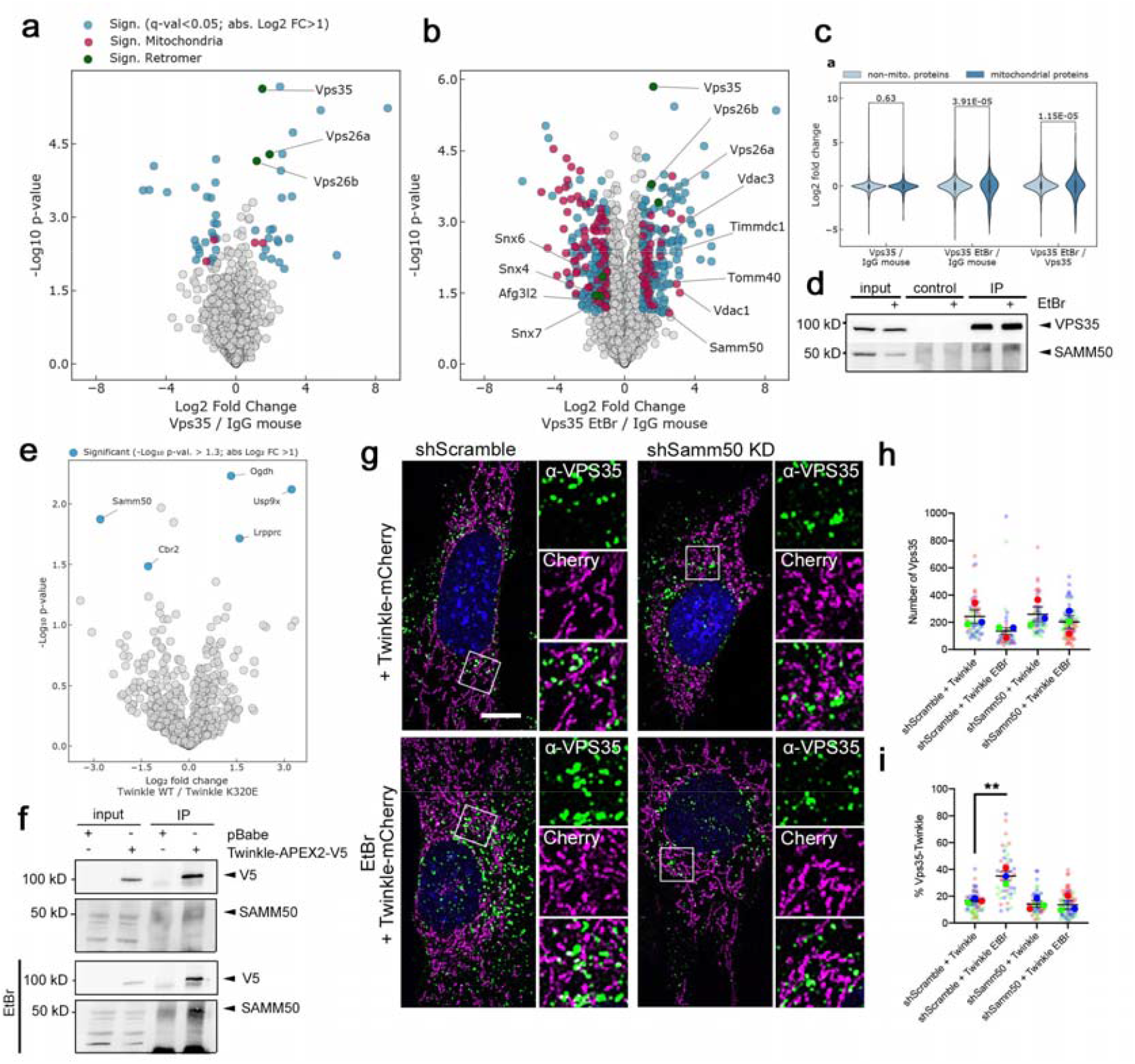
SAMM50 is a mitochondrial receptor for VPS35 recruitment into mitochondrial subcompartments containing nucleoids. (**a**) Volcano plot showing proteins enriched after immunoprecipitation of VPS35 in MEFs. Only differentially enriched proteins are highlighted (adj. p-value <0.05 and FC >2 or <-2). Blue, non-mitochondrial proteins; Red, mitochondrial proteins; Green, retromer proteins (**b**) Comparison of VPS35 IP profiles of cells grown in normal medium or 1 week treated with 50ng/ml EtBr. n=3. (**c**) Violin Plots showing enrichment of mitochondrial proteins upon pull down of VPS35 in presence of EtBr. Numbers indicate p-value. Students’
s T-test. (**d**) Co-Immunoprecipitation of VPS35 and SAMM50 in steady state and EtBr treated cells. IgG from mouse was used as a control. (**e**) Comparison of interactome profiles of Twinkle and K320E. n=3. (**f**) co-Immunoprecipitation of Twinkle and SAMM50 in steady state and in cells treated with EtBr for 1 week. (**g**) Immunofluorescence of control and Samm50 KD MEFs transduced with Twinkle-mCherry constructs and labelled with α-VPS35 in basal and 7 days treated with 50ng/ml EtBr. Bar, 10µm. (**h, i**) Quantification of VPS35 particles and VPS35 in contact with Twinkle. n=3, >20 cells per replicate. ANOVA, Tukey multiple comparison. **, p<0.01. Mean ± SEM

Thus, we hypothesize that SAMM50, a protein located in the mitochondrial outer membrane, could serve as a platform to recruit VPS35 and eliminate mitochondrial subcompartments containing damaged mtDNA. Consequently, we generated SAMM50 shRNA KD clones and expressed Twinkle-mCherry (Fig. 7g; Fig. S8c, d). EtBr does not affect the overall population of VPS35 but enhances recruitment to Twinkle (Fig. 7g-i). However, the percentage of VPS35-Twinkle contacts were reduced in absence of SAMM50 (Fig. 7i). To decipher if VPS35 recruitment shows Twinkle specificity, we examined VPS35 colocalization with LRPPRC, a native mitochondrial matrix protein (Fig. S8e). Again, the overall number of VPS35 particles was unchanged by EtBr treatment (Fig. S8f) and in this case, we observed that VPS35 contacts with LRPPRC were not increased and therefore, there was no effect following Samm50 downregulation (Fig. S8g).

In summary, our data thus demonstrate that VPS35 is required for mitochondrial quality control in the presence of mtDNA alterations. We show that a mechanism to specifically remove mtDNA is activated when mtDNA damage is induced both genetically or chemically by K320E or by EtBr, respectively. ATAD3 is required for specific membrane localization of nucleoids, SAMM50 confers nucleoid specificity and VPS35 is involved in fine-tuning of this selective process, removing specific parts of mitochondria containing affected nucleoids, thus avoiding the activation of acute mitophagy which would affect the functional mitochondrial pool.

### Rapamycin eliminates mtDNA alterations without affecting copy number

Since our data highlight that autophagy, together with a specific nucleoid extraction mechanism, plays an important role in maintaining mtDNA fitness *in vitro*, we aimed to test whether this is also relevant *in vivo*. We have previously shown that expression of K320E in skeletal muscle leads to the accumulation of mtDNA alterations, unfortunately only in very old animals, making this model not very convenient ^22^ (Fig. 1). However, expression of K320E in muscle satellite cells (Pax7-Cre^ERT^; K320E^msc^), followed by cardiotoxin-induced muscle damage and one week of regeneration, shows a rapid accumulation of mtDNA alterations, leading to newly generated, cytochrome-c-oxidase (COX) negative fibers (stained blue, Fig. S8a, b), while mtDNA copy number remained stable (Fig. S8c). In several mitochondrial disease models, it has been shown that rapamycin, an activator of autophagy through the specific inhibition of mTORC1, is able to revert mitochondrial dysfunction and ameliorate disease progression. This probably occurs by stimulating mitochondrial turnover mechanisms ^35^, and, *in vitro* it was also shown to direct selection against mtDNA mutations ^36^. Interestingly, genetic induction of autophagy was shown to reduce mtDNA deletions in *Drosophila* ^37^. Thus, to test if activation of autophagy can purify mtDNA alterations also in mammals *in vivo*, we used this muscle regeneration paradigm in combination with rapamycin treatment. Regenerated muscles from vehicle treated K320E^msc^ mice showed a prominent accumulation of COX negative fibers indicating mitochondrial dysfunction (Fig. 8a). In contrast, mice treated with rapamycin showed much less COX deficient cells in the regenerated area (Fig. 8a-c). Consistently, qPCR analysis revealed that mtDNA copy number remained unchanged (Fig. 8d), while mtDNA alterations were absent and thus had been purified (Fig. 8e). We noticed that COX staining was much lighter in rapamycin treated animals, suggesting a change in mitochondrial OXPHOS activity. Vehicle treated mice showed a predominant accumulation of glycolytic fiber type 2b in both wt and K320E^msc^ mice (Fig. S9d), while in rapamycin treated wt mice, fiber type staining showed a predominant shift towards mitochondria rich type I fibers (Fig. S9e). Interestingly, in K320E^msc^ mice regenerated fibers showed a mixed myosin heavy chain pattern after one week of regeneration. These data indicate that rapamycin can be used as a modulator of mitochondrial turnover which specifically purifies mutated mtDNA species, thus ameliorating mitochondrial dysfunction, albeit changing muscle fiber type composition.

**Figure 8.**
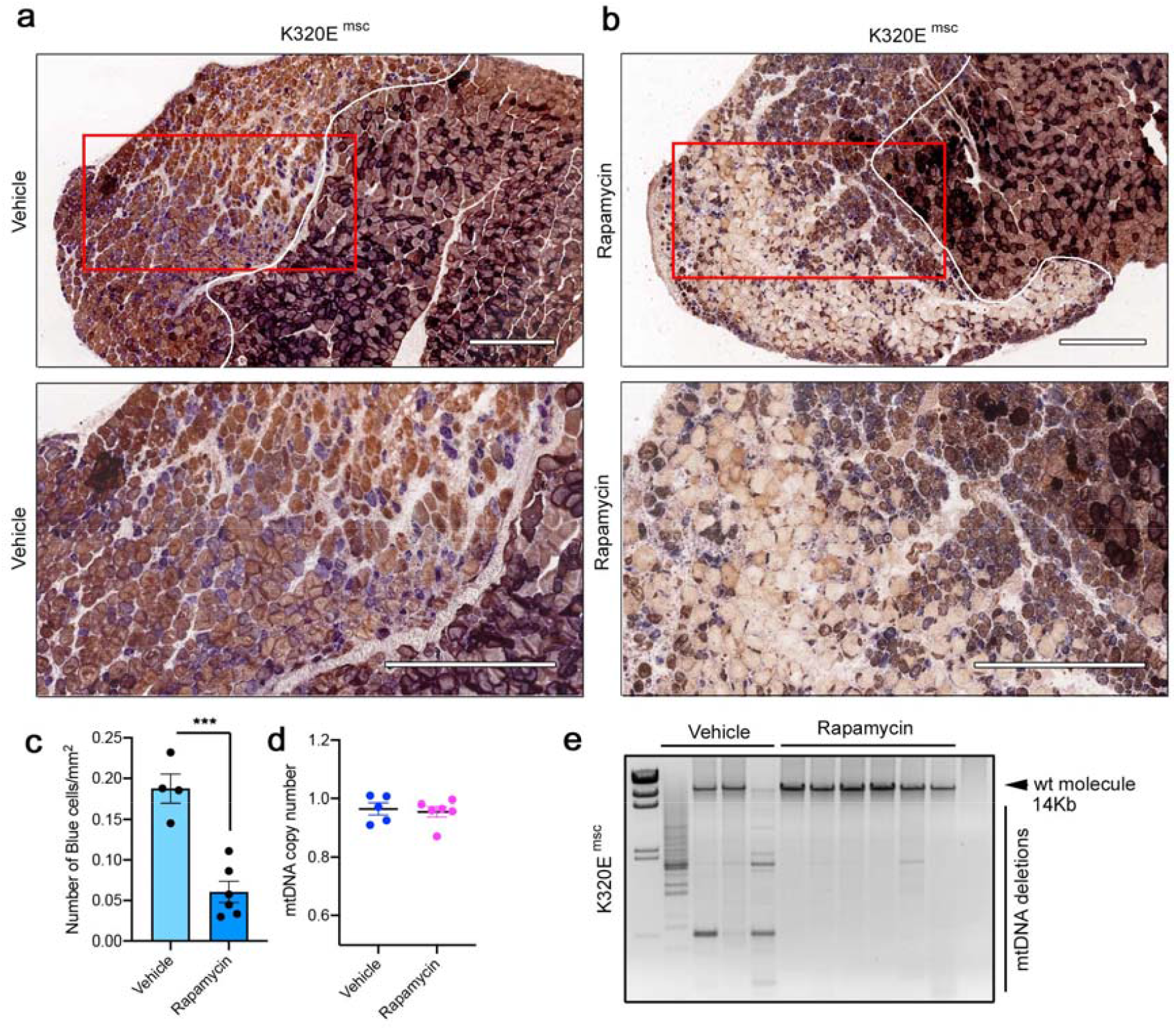
Rapamycin eliminates mtDNA deletions without affecting copy number *in vivo*. COX-SDH staining of regenerated TA muscle from Pax7-K320E mice (K320E^msc^). After cardiotoxin induced injury, mice were injected for five days either with (**a**) vehicle or (**b**) 2 mg/Kg Rapamycin. Bar, 500µm. (**c**) Quantification of COX negative cells (blue) in the injured area. (**d**) mtDNA quantification by qPCR or (**e**) Long-range PCR in regenerated muscle from K320E^msc^ mice treated with vehicle or with Rapamycin. n=5-6 mice per condition, genotype and treatment. (**c** and **d**) Unpaired Students’
s T-test. Mean ± SEM.

## Discussion

Autophagy and specifically mitophagy and its variants are well-established pathways for mitochondrial turnover, which is essential to maintain mitochondrial fitness ^4^. Loss of mitochondrial quality control mechanisms, either by specific mutations of key players or by reduced autophagy activity, strikingly correlates with the acquirement of mtDNA mutations ^38^. However, an exacerbated activation of mitophagy may lead to a serious reduction of the mitochondrial pool, thus affecting cellular energy supply ^39^. Therefore, the fine-tuned regulation of mitochondrial quality control mechanisms is essential to maintain cellular energy homeostasis.

Although toxic substances can cause mitochondrial damage, the most prevalent reason is the accumulation of alterations in mtDNA due to replication errors. Mutations in genes encoding proteins involved in mtDNA replication, like the helicase Twinkle, and in mtDNA maintenance cause mutations leading to mitochondrial diseases, with brain and skeletal muscle being regularly affected. In addition, somatic mutations in mtDNA accumulate during the normal aging process in many organs in humans ^3^, leading to a tissue mosaic where few cells with mitochondrial dysfunction, caused by high mutation loads, are embedded in normal tissue ^40^.

In general, tissues most depending on mitochondrial function are most severely affected when carrying mtDNA mutations. Paradoxically, we found that expression of the dominant negative K320E mutation of Twinkle in extraocular muscle shows a remarkable differential vulnerability of muscle fiber types, with mitochondrial dysfunction especially affecting fibers with a glycolytic metabolism ^22^. In agreement with these results, we found less mtDNA alterations in aged SOL, a muscle rich in type I fibers which mostly rely on mitochondrial ATP production, compared to the TA mostly composed of fast twitch, glycolytic fibers (Fig. 1). Noteworthy, different muscles rich in oxidative vs. glycolytic fibers show notable differences in the expression of genes involved in mitochondrial dynamics ^41^, making oxidative muscles more resistant to ageing related dysfunction ^42^. In fact, our data shows that SOL expressing K320E already has an increased flux in steady state but this was not related to increased mitochondrial turnover. Nevertheless, mitophagy, understood as the specific removal of the entire damaged organelle, does not provide the required selectivity to remove only mutated mtDNA. Hence, the existence of a specific turnover mechanism has been postulated, but not been proven yet ^43^.

In contrast to terminally differentiated muscle of aged mice, proliferating cells in culture did not accumulate mtDNA alterations upon expression of K320E, instead this led to mtDNA depletion. Both K320E expression or mild treatment with EtBr induced the accumulation of mtDNA damage. EtBr intercalates between base pairs and slows down mtDNA replication. It is accepted that EtBr, when used at low concentration, also induces frame shift mutations and deletions^44^ and we demonstrated that it also triggers oxidative damage in the mtDNA. Our data show that mtDNA depletion induced by damage is caused by a specific exacerbated mtDNA turnover which is Atg5 dependent and thus requires autophagosome formation and lysosomal degradation.

Interestingly, K320E localizes preferentially in specific mitochondrial regions. K320E localization does not overlap completely with the mitochondrial outer membrane protein TOM20 and is present in areas that are also not stained by Mitotracker Green, demonstrating that nucleoids containing Twinkle can be specifically localized in a unique mitochondrial subcompartment. Indeed, mtDNA has been shown to attach preferentially to cholesterol-rich membrane structures and Twinkle, as a nucleoid protein, has been found to be enriched in these areas as well ^45^. The onion-like structures we observed in our cell model have been detected in the muscle of patients with mitochondrial myopathy caused by mtDNA mutations ^46^ and very recently also in ρ0-cells lacking mtDNA ^47^. Here we show that mtDNA damage induces cristae remodelling in poles of the mitochondrial network and that these structures containing Twinkle are delivered to lysosomal compartments, which are therefore specifically involved in mtDNA removal.

Three proteins present in three mitochondrial compartments are responsible for mtDNA distribution and selective turnover: Twinkle, in the mitochondrial matrix; ATAD3, in the mitochondrial inner membrane and SAMM50, in the mitochondrial outer membrane. Twinkle arises as the link between nucleoids and the inner membrane through interaction with ATAD3, a protein controlling several aspects of mitochondrial membrane dynamics ^48^. Interestingly, the human ortholog ATAD3B has been recently found to be a mitophagy receptor for mtDNA damage induced by oxidative stress ^49^. In the other side, SAMM50, which resides in the mitochondrial outer membrane, interacts with the MICOS complex, and organizes membrane architecture ^50^. Interestingly, both ATAD3 and SAMM50 have been described as regulators of Pink1-Parkin as well ^34,51^, thus providing the link between nucleoid localization and specific degradation of mtDNA.

Furthermore, we show that upon mtDNA damage, VPS35, an endosomal protein involved in mitochondrial quality control, increases contacts to mitochondrial subcompartments containing Twinkle. Interestingly, VPS35 deficient cells showed a persistent activation of mitophagy with mtDNA depletion. The fact that *Samm50* depletion induces mitophagy, but excludes mtDNA degradation ^34^, suggests that SAMM50 confers the required selectivity and VPS35 the specificity to remove only fragments containing mtDNA.

Removal of mitochondrial fragments has been shown to take part through a specialized pathway called Mitochondrial Derived Vesicles (MDVs) ^7^. Currently, only a few proteins have been found to determine cargo and vesicle fate. While MAPL generates vesicles which divert oxidized cargo to peroxisomes ^32^, Tollip, an endosomal organizer, synchronizes Parkin-dependent MDVs directed to the lysosomal compartment ^8^. Canonical MDVs are generated in the mitochondria in an Atg5 and LC3 independent manner ^32^, however, mtDNA damage by K320E expression showed accumulation of LC3 autophagosomes and mtDNA depletion which is Atg5 dependent. Additionally, electron microscopic pictures of our cells never showed vesicles resembling MDVs, which are much smaller than the mitochondrial extrusions and autolysosomes containing Twinkle we observed. Furthermore, we unequivocally showed that Twinkle particles are directed to the lysosomal compartment and that VPS35 associates with mitochondria upon mtDNA damage. We also showed that VPS35 recruitment is SAMM50 dependent and, upon mtDNA damage, such recruitment is specific to Twinkle containing regions. Upon EtBr damage, we confirmed that VPS35 precipitates with mitochondrial outer membrane proteins, such as SAMM50, TOMM40, TIMMDC1 and VDAC1 and VDAC3 and it is in this environment when Twinkle, a mitochondrial matrix protein, interacts with SAMM50. mtDNA removal exemplifies a specialized mitophagy trail. Delivery of specific mitochondrial cargo to quality control pathways through VPS35 was previously shown to be highly regulated. If the selected cargo is diverted into peroxisomes, VPS35 associates with MDVs by interacting with MAPL in a Drp1 independent manner ^10^. However, if the MDVs are directed to lysosomes the association is dependent on Drp1 ^52^. On the other hand, SAMM50 modifies mitochondrial structure by interacting with Drp1 ^53^ and with the Pink1-Parkin machinery for selective mitophagy ^34^. Hence, SAMM50 works as a platform where mitochondrial fission, membrane architecture, and mitophagy components orchestrate membrane protrusions and facilitate VPS35 recruitment. Very recently, SAMM50 has been linked also to piecemeal mitophagy, independent of MDVs, through direct interaction with the p62 adaptor ^54^. Hence, we conclude that specific mtDNA degradation does not follow the MDV pathway, despite the fact that they share some components, and that it is more likely that mitochondrial protrusions containing nucleoids are engulfed in autophagosomes in a specialized mitophagy pathway.

VPS35, which has been extensively linked to neurological diseases such as Parkinson’s and Alzheimer disease, appears as a regulator of mtDNA quality control necessary to maintain mitochondrial intactness. Modulation of *VPS35* expression has been evaluated as a potential approach against Parkinson’s disease ^55^ and, in *Drosophila, Vps35* overexpression can rescue an LRRK2-induced Parkinson’
ss phenotype ^56^. iPSC derived neurons from *LRKK2*-PD patients showed accumulation of mtDNA damage ^57^. Nonetheless, the ability of VPS35 to eliminate such molecules in a human disease related model needs to be further explored as a potential therapeutic strategy.

Enrichment of Spg7 upon VPS35 IP in EtBr treated cells also suggests the involvement of mitochondrial proteases. Mitochondrial matrix proteases (mAAAs) represent a group of enzymes related to mitochondrial quality control and mitochondrial membrane remodeling upon proteolytic cleavage of Opa1 and Oma1. Spg7 has been found to copurify with Prohibitin participating in the formation of the permeability transition pore ^58,59^. Mutations in the *SPG7* gene are the cause for Hereditary Spastic Paraplegia Type 7 but also a Progressive External Optalmoplegia-like syndrome with accumulation of mtDNA deletions ^60^. Hence, it is tempting to propose that Spg7 works as a regulator of mtDNA turnover as well. However, further studies are needed to reveal its specific role and regulation of the process.

Finally, our data demonstrate that specific removal of mtDNA is linked to lysosomal activity. Lysosomal degradation of bulk autophagy is regulated by the serine-threonine protein kinase mTORC1 which resides on the lysosomal surface. mtDNA replication defects activate mTORC1 and the integrated mitochondrial stress response in a cascade of effects with wide downstream consequences ^61^. It is well known that mTORC1 activation inhibits autophagy by influencing both the formation of autophagosomes but also lysosomal acidification ^62^. In order to prove the selectivity of this process *in vivo*, we used a mouse model where mtDNA alterations rapidly accumulate and indeed, activation of the mTORC1 pathway by rapamycin was able to eliminate abnormal mtDNA molecules and thus reduces the accumulation of cells with mitochondrial dysfunction. Rapamycin has been described as a potential treatment against mitochondrial diseases ^18,63^, however, since we observed a fiber-type shift, mTORC1 inhibition might activate other signalling pathways with undesirable effects. Nonetheless, by using Rapamycin we demonstrated that elimination of mutant mtDNA without affecting total mtDNA copy number is possible.

In conclusion, we unveil a new complex mechanism with physiological relevance for mitochondrial fitness. Twinkle mediates nucleoid binding to the mitochondrial inner membrane through ATAD3 interaction, which is responsible for nucleoid organization. SAMM50 provides the required specificity to eliminate mtDNA while VPS35 supplies the selectivity. Interestingly, mutations in *TWNK, ATAD3A* and *VPS35* have been linked to several severe mitochondrial diseases having in common mtDNA instability ^64-66^, therefore representing a cluster of proteins involved in specific mtDNA turnover. Fine tuning the activity of such proteins could be used as a therapeutic strategy against mtDNA related diseases, either inherited, acquired or due to normal ageing.

## Methods

### *In vivo* experimental approaches

K320E transgenic mice were generated by crossing R26-K320EloxP/+ mice (point mutation K320E; Rosa26-Stop-construct; downstream EGFP) with mice expressing Cre recombinase under the control of the skeletal muscle-specific MLC1f-promoter or satellite cells Pax7-Cre^ERT^. All mice used for experiments were housed in a standard animal facility maintained at 23°C, 12:12h light-dark cycle, with free access to water and standard rodent chow. Autophagy flux was tested by intraperitoneal injection of 50mg/kg chloroquine 4h prior euthanasia. All procedures and experimentation with mice were performed according to protocols approved by the local authority (LANUV, Landesamt fu□r Natur, Umwelt und Verbraucherschutz NRW, approval number: 2019-A090). Activation of Pax7-Cre^ERT^ promotor was performed by injecting daily, for 5 days, intraperitoneal 2mg tamoxifen dissolved in mygliol. For muscle regeneration experiments, 2 days after the last tamoxifen injection, mice were anesthetized with 2%Xylazin, 10% Ketamine in NaCl 0.9% and 10μM Cardiotoxin (*Naja Pallida*, Latoxan) was injected inside the fascia. After 2 days of rest, 2mg/kg rapamycin dissolved in mygliol was injected intraperitoneally daily for 5 days. The day after the last injection, mice were treated as explained below.

### Molecular biology and vector generation

*Twnk* ORF was amplified from plasmids pJet2-Twinkle and pROSA-K320E ^67^ and cloned in pmCherry-N1. For retroviral vector generation, Twinkle and Twinkle-mCherry, ORF was amplified and subcloned into pLenti-CMV Puro DEST (Addgene #17452). All ORFs were subcloned into pBabe-Puro vector, kindly provided by Dr. Bernhard Schermer, and verified by Sanger sequencing. pLX304-TWINKLE-APEX2 vector was kindly provided by Dr. Alice Ting. To generate *Mus musculus Twnk* vectors, *Homo sapines TWNK* was replaced by *Twnk* ORF and subcloned into pBABE-Puro vector. Mitochondrial matrix APEX2 control vector (Addgene #72480) was also subcloned into pBABE-Puro. For generation of Tfam-GFP, total RNA from mouse liver was isolated and converted into cDNA using TRIzol (Thermo Fisher) and RevertAid First Strand cDNA Synthesis Kit (Thermo Fisher) using PolyT as feeder. *Tfam* cDNA was cloned into pEGFP-N1. Atad3 KD and Samm50 KD clones were generated by transducing MEFs with the vector pMKO1-GFP (Addgene #10676) containing a shRNA (Table M1). Empty vector and a scramble containing vector were used as a control. For CRISPR Cas9 KO generation two gRNA directed to exon 4 and exon 5 (Table M1) were cloned in pSpCas9 (BB)-2A-Puro V2.0 (Addgene #62988).

### Generation and culture of cell lines

C2C12 cell line was purchased from ATTC and grown in DMEM 4.5 g/L Glucose + GlutaMax, 20% FBS, 1x Pen/Strep. Immortalized mouse embryonic fibroblast Atg5WT and Atg5KO MEFs ^68^ and immortalized MEFs line were maintained as described ^69^. Stable cell lines were generated by transducing C2C12 cells or MEFs with pBABE-Puro retroviruses. Briefly, 2.5×10^6^ HEK293 cells were plated in 10cm^2^ dish transfected with pCL-ECO (5 μg) and pBABE-Puro (10 μg) vectors using PEI (40 μl). After 48h and 72h, medium containing viruses was harvested, filtered through 45 μm, mixed with 8μg/ml Polybrene and added to 250.000 cells previously plated. 48h post transduction, positive clones were selected by adding Puromycin 2.5 μg/ml to the medium in C2C12 and 5μg/ml to MEFs, which was maintained during all the experiments. shRNA clones were generated by transducing MEFs with pMKO.1-GFP vectors. Prior to all experiments, transduction rate was verified to be higher than 99% of GFP expressing cells by Flow cytometry. For generation of Vps35 CRISPR Cas9 KO clones, MEFs were transiently transfected with the vector containing gRNA and selected with 3 µg/ml puromycin for 4 days. Single clones were plated independently using cloning cylinders, analyzed by western blotting. Genomic DNA was isolated from VPS35-negative cells and genomic DNA modification was verified by Sanger sequencing. Exon 4 and Exon 5 were amplified (Table M1) and cloned using pJET1.2 cloning kit (Thermo Fisher) before sequencing.

Transient transfection was achieved transfecting the corresponding plasmids using Lipofectamine 3000 following manufacturer instructions. Plasmids use in this work were: LC3-GFP (Addgene #21073), Lamp1-GFP (Addgene #34831), Fis1p-Cherry-GFP ^70^, LC3-Fis1p-Cherry (kindly provided by Dr. Terje Johansen) and MAPL-GFP from CECAD Imaging Facility ^32^.

For mtDNA depletion experiments, cells were maintained as described before for 7 days but media was supplemented with 50 ng/ml Ethidium and 50 μg/ml Uridine. Lysosomal function was blocked using 10 μM Chloroquine for 24h.

### mtDNA amplification

Total DNA was isolated using DNeasy Blood & Tissue Kit (Qiagen) according to manufacturer’
ss instruction. 25 ng of total DNA was used for analysis of threshold amplification differences between mtDNA and nuclear DNA (delta C(t) method with specific primers (Table M1). Long range PCR was used to screen for the presence of mtDNA alterations. 14 Kb of mtDNA was amplified using Rabbit Bioscience Long Range kit with oligos described in table M1.

### Western blot and co-immunoprecipitation

Cells pellets were lysed with RIPA-buffer (150 mM NaCl, 1% Triton-X1000, 0.1% SDS, 50 mM Tris-HCl pH 8, 0.5% Na-deoxycholate) containing protease inhibitor (Roche) and protein concentration measured using the Bradford assay. Proteins were transferred after electrophoresis to a PVDF-membrane previously activated with methanol. Membranes were blocked (5% milk in TBS-0.1% Tween-20) and incubated overnight with primary antibodies. Antibodies used in this work are: monoclonal α-V5 (Abcam), polyclonal α-V5 (Thermo Scientific), polyclonal α-TOM20 and monoclonal α-VPS35 (Santa Cruz), monoclonal and polyclonal α-SAMM50 (Abnova and Abcam respectively), polyclonal α-ATAD3, polyclonal α-LC3 and polyclonal α-p62 (Proteintech), polyclonal GAPDH (Novus Biologicals). Secondary goat anti-mouse, goat anti-rabbit and goat anti-chicken HRP (Jackson Laboratory). Images were acquired using the ECL Advanced Chemiluminescence kit (GE Healthcare Life Sciences ®, UK) according to manufacturer’s protocols and visualized using a LAS500 CCD camera.

For immunoprecipitation, cells expressing Twinkle-APEX2-V5 were pelleted and solubilized in IP Buffer (500 mM HEPES KOH pH 7.2, 150 mM NaCl, 1 mM MgCl_2_, 1% Triton-X1000 and Protease inhibitor (Roche). 500 µg of total protein extract were used to IP with either 2.5 µg V5 rabbit polyclonal antibody or VPS35 mouse monoclonal over night at 4°C and recovered after incubating for 6h at 4°C in a rotator with equilibrated Agarose Protein-G beads (Abcam). Immunoprecipitation followed by MS analysis was performed as described but using magnetic Protein G beads (Thermo Scientific). Samples were washed 5 times with washing buffer (10 mM HEPES KOH pH 7.2, 150 mM NaCl, 1 mM MgCl_2_, 0.2% Triton-X1000) and once with PBS. Prior to analysis by MS, samples were washed 3x with ammonium-bicarbonate (ABC) buffer, denatured with 50 µl of urea buffer (6 M urea, 2 M thiourea) and followed by disulfide-bridge reduction using dithiothreitol (DTT) at a final concentration of 5 mM for 1 hour at room temperature. To alkylate oxidized cysteines, 2-Iodoacetamide (IAA) was added to the samples until a concentration of 40 mM was reached and incubated for 30 min in the dark. Lys-C was added in a ratio of 1:100 (0.1 µg enzyme for 10 µg protein) and incubated for 2-3 hours. Samples were finally diluted with ABC buffer to reach 2M urea concentration. Protein digestion was performed overnight with trypsin 1:100. Samples were acidified with 1% formic acid and desalted using a modified version of the previously described Stop and Go extraction tip (StageTip) protocol ^71^.

### Pulsed SILAC labeling in mice and in-solution digestion

For pulsed SILAC labeling mice, 30-40 weeks old mice for K320E; Mlc1 line (C57BL/6J) were fed a ^13^C_6_-lysine (Lys6)-containing mouse diet (Silantes) for 14 days to monitor newly synthesized proteins by comparing the incorporation of Lys-6 with the naturally occurring Lys-0 ^72^. Mice were sacrificed at the end of day 14 and tissues dissected and snap-frozen in liquid nitrogen. Samples were grinded and proteins extracted and denatured by the addition of 4% SDS in PBS. To remove residual SDS proteins were precipitated overnight in 4x ice-cold acetone (v:v). On the next day after centrifugation at 16,000 g for 10 min the protein pellets were dissolved in urea buffer (6 M urea / 2 M thiourea). The following protein digestion was performed as described previously but instead of overnight tryptic digestion proteins were only digested with Lys-C (1:100 enzyme-to-protein ratio) for both pre-digestion (2h at RT) and overnight digestion after dilution of urea using ABC buffer.

### Liquid chromatography – mass spectrometric analysis

Both affinity-enriched and pulsed SILAC proteomics samples were analyzed in positive mode using data-dependent acquisition (DDA) either by an Easy-nLC 1000 – Q Exactive Plus or an Easy-nLC 1200 – Orbitrap Eclipse tribrid system (all Thermo Fisher). On-line chromatography was directly coupled to the mass spectrometric systems using a nanoelectrospray ionization source. Peptides were separated by reversed-phase chromatography with a binary buffer system of buffer A (0.1% formic acid in water) and buffer B (0.1% formic acid in 80% acetonitrile) using a 60 min chromatographic gradient for IP samples and 120 min for pulsed SILAC samples. Separation was performed on a 50 cm long in-house packed analytical column filled with 1.9 µM C18-AQ Reprosil Pur beads (Dr. Maisch). Using the 60 min chromatographic gradient peptide separation based on their hydrophobicity was performed by linearly increasing the amount of buffer B from initial 13% to 48% over 35 min followed by an increase of B to 95% for 10 min. The column was washed for 5 min and initial column conditions were achieved by equilibrating the column for 10 min at 7% B. Full MS spectra (300 to 1750 m/z) were acquired with a resolution of 70,000, a maximum injection time of 20 ms and an AGC target of 3e6. The top 10 most abundant peptide ions were isolated (1.8 m/z isolation windows) for subsequent HCD fragmentation (NCE = 28) and MS/MS recording at a resolution of 35,000, a maximum injection time of 120 ms and an AGC target of 5e5. Peptide ions selected for fragmentation were dynamically excluded for 20 seconds.

Using the 120 min chromatographic gradient peptides were separated by linearly increasing B from initial 4% to 25% over 96 min followed by an increase of B to 55% over 14 min. After a steep increase of B to 95% over 2 min the analytical column was washed for 8 min at 95% B. Full MS spectra (375 to 1500 m/z) were acquired with a resolution of 60,000, a dynamic injection time and an automated AGC target. The top 20 most abundant peptide ions (charge state 2 – 7) were isolated (1.2 m/z isolation windows) for subsequent HCD fragmentation (NCE = 30) and MS/MS recording at a resolution of 15,000, a maximum injection time of 22 ms and an automated AGC target. Peptide ions selected for fragmentation were dynamically excluded for 60 seconds.

### Data processing and analysis

The mass spectrometry proteomics data have been deposited to the ProteomeXchange Consortium (http://proteomecentral.proteomexchange.org) via the PRIDE partner repository ^73^ with the dataset identifier PXD023939. All recorded RAW files were processed with the MaxQuant software suite (1.5.3.8 for IP data, 1.6.14 for pSILAC) ^74^. For peptide identification and scoring MS/MS spectra were matched against the mouse Uniprot database (downloaded 08/15/2019) using the Andromeda search algorithm ^75^. For the affinity-enriched samples multiplicity was set to one and trypsin/P was selected as digestive enzyme. Carbamidomethylation was set as a fixed modification and methionine oxidation or N-terminal acetylation was selected as variable modification. Peptides were identified with a minimum amino acid length of seven and a false-discovery rate (FDR) cut-off of 1% on the peptide level. Proteins were identified with FDR < 1% using unique and razor peptides for quantification. Label-free quantification was performed using the standard settings of the maxLFQ algorithm. Match between runs was activated. pSILAC data were analyzed with the same settings with some modifications. Multiplicity was set to two with Lys6 as heavy isotope label. Lys-C/P was selected as digestive peptide and LFQ quantification was deactivated.

Statistical analysis and visualization were done with the Perseus (1.6.5) and InstantClue software suits ^76,77^. LFQ intensities of the IP samples were log_2_-transformed and filtered for proteins identified in all replicates of at least one condition. Missing values were imputed by random drawing of values from 1.8 standard deviations (SD) downshifted, 0.3 SD broad normal distribution to simulate the lower detection limits of the mass spectrometer. To evaluate principal components responsible for the variances between samples we performed a principal component analysis. Further, we performed a two-sided Student’s *t*-test to identify significantly regulated proteins. We proceeded similarly for the pulsed SILAC data but used the heavy-to-light (H/L) ratios for statistical testing. We used the same filtering-criteria but did not imputed missing values. We performed a 1D annotation enrichment to identify enriched categorical terms in the different conditions ^78^.

### Histology, Immunofluorescence and Microscopy

For tissue histology, mice were sacrificed by cervical dislocation, muscles dissected, mounted in cork with OCT (Tissue-Tek), snap frozen in isopentane and stored at -80°C till needed. 10 µm thick sections covering the injured area were produced using a cryostat maintained at -20°C (Leica CM 3050s, Techno-med). To assess the integrity of mitochondrial function the sections were sequentially stained for COX and SDH activities. Frozen sections were incubated 20 mins at 37°C in COX solution (20 mg/ml catalase, 74 mg/ml sucrose, 2 mg/ml cytochrome c, and 1mg/ml DAB in 50mM Na_2_HPO_4_ pH 7.4). After 3 PBS washes, sections were then incubated for 30 min at 37°C in SDH staining solution (2 mg/ml NBT, 0.2M Sodium succinate, 50mM MgCl_2_, 50mM Tris-HCl, pH 7.4), washed 3 times with miliQ water, and mounted in Glycerol gelatin medium (Sigma).

For immunofluorescence of autophagy markers LC3 and p62 and mitochondrial TOM20, mice were perfused with PFA 4% in PBS prior to muscle collection. Samples were equilibrated in 15% sucrose for 6h and 30% sucrose overnight before frozen in OCT medium. For LC3 IF, samples were preincubated with 0.1% SDS for 5min. Antibody specificity was determined in muscle sections from LC3-GFP transgenic mice (kindly provided by Dr. Evangelos Kondilis). Cryosections were blocked for 1h with 1% Western blocking reagent (Roche) containing 0.1% Triton in PBST, antibodies incubated overnight at 4°C and secondary antibodies at room temperature for 1h in blocking buffer. Samples were mounted in Fluoromount G containing DAPI. Fiber type staining was performed as described previously ^26^. Images were obtained with Leica SP8 with 63x/1.40 oil PL Apo objective.

For *in vitro* analysis, cells were fixed in 4% PFA/PBS, permeabilized with PBS-0.2% Triton-X1000 for 30 min and blocked for 1 hour at RT in blocking buffer (5% fat free milk powder, 10% FBS, 1% BSA, 0.1% Triton-X100 in PBS). Primary antibodies were incubated in blocking buffer over night at 4°C and secondary antibodies for 1 hour at RT. mtDNA replication rate was determined by pulse BrdU labelling. Briefly, cells were incubated with 20 μM BrdU (Sigma) for 6h, fixed with 4% PFA/PBS for 30 minutes and directly permeabilize with 05% Triton X-100 on ice for 5 min. To allow access to mtDNA, cells were incubated with HCl 2N for another 30 min prior to immunofluorescence. Coverslips were mounted using DAPI-Fluoromount G. Images were acquired using a spinning-disk confocal microscope (Ultra View VoX; PerkinElmer) with a Plan-Apochromat total internal reflection fluorescence 60Å∼/1.49 NA oil DIC objective, Leica SP5 microscope controlled by Las AF 3 with 2.5x extra magnification and Leica SP8 with 63x/1.40 oil PL Apo objective. Microscopy Live imaging was performed in spinning-disk confocal microscope (Ultra View VoX; PerkinElmer). Cells were seeded onto glass plates and loaded with 500nM Mitotracker green for 30 minutes. Videos were recorded taking 1 picture/min at 37°C and 5% CO_2_.

Antibodies used for immunofluorescence were: rabbit polyclonal α-V5 (Thermo); rabbit polyclonal α-TOM20, mouse monoclonal α-Vsp35 and mouse monoclonal α-8-OHdG (Santa Cruz); goat polyclonal α-Vsp35 and mouse monoclonal α-dsDNA (Abcam); rabbit polyclonal α-ATAD3, rabbit polyclonal α-LC3, rabbit polyclonal α-p62, rabbit polyclonal LRPPRC (Proteintech); and monoclonal α-BrdU (BD Bioscience). Fluorescence secondary antibodies goat α-mouse, α-rabbit Alexa Fluor-488, 555 and 647 and rabbit α-goat-647 were used accordingly to the primary antibodies. Additionally, α-mouse IgM Alexa Fluor 488, α-mouse IgG Alexa Fluor 555 and α-mouse IgG2b Alexa Fluor 647 were used for fiber type triple staining.

### Electron microscopy

For electron microscope cells were grown on small discs of aclar foil and fixed for 1 h in 2% Glutaraldehyde with 2 %Sucrose in HEPES buffer pH 7,4. After washing two times with 0.1M Cacodylate buffer, free aldehyde groups were quenched with 0.1M Glycin in 0.1M Cacodylate buffer for two times 20 min. After a short wash with 0.1M Cacodylate buffer, cells were incubated in 0.5mg/ml Diaminobenzidine in 0.1M Cacodylate buffer and, after 10 min, a final concentration of 0.03% H_2_O_2_ added and incubated for 30min. Finally, cells were washed three times with 0.1M Cacodylate buffer and incubated with 1% Osmiumtetroxid and 1.5% Potassium hexacyanoferrat for 30 min at 4°C. After 3×5min wash with ddH2O, samples were dehydrated using ascending ethanol series (50%, 70%, 90%, 100%) for 5 min each and infiltrated with a mixture of 50% Epon/ethanol overnight at 4°C and with pure Epon for two times 2 h. Samples were embedded into TAAB capsules and cured for 48 h at 60°C.

For electron tomography Ultrathin sections of 200 nm were cut using an ultramicrotome (Leica, UC7) and incubated with 10 nm protein A gold (CMC, Utrecht) diluted 1:25 in ddH20. Sections were stained with 2% Uranyl acetate for 20 min and Reynolds lead citrate solution for 3 min. Images and Tilt series for 1nm thickness were acquired from -65° to 65° with 1° increment on a JEM-2100 Plus Transmission Electron Microscope (JEOL) operating at 200kV equipped with a OneView 4K 32 bit (Gatan) using SerialEM (Mastronarde, 2005). Reconstruction was done using Imod (Kremer et al.,1996).

### Image analysis

All image analysis was performed in FIJI (NIH, Bethesda). LC3 and p62 puncta quantification were performed with the “counting cells” internal plugin for particles bigger than 2 pixels to exclude background. mtDNA foci quantification, VPS35-Twinkle analysis, 8-OHdG and BrdU analysis were performed with a self-created macro. Briefly, threshold was set for the different channels. Nuclear signal was selected and removed for the analysis. The signal corresponding to the mitochondrial network was selected and only the particles from the other channel bigger than 1 pixel inside the mitochondrial network were considered to the analysis. For VPS35, the minimum size was determined to be 3 pixels. After thresholding, all VPS35 particles were summarized and the ones in contact with Twinkle or LRPPRC were used to get a percentage. Manders’
s coefficient was obtained using JaCOP plugin. (Bolte & Cordelieres, 2006).

**Table M1.**
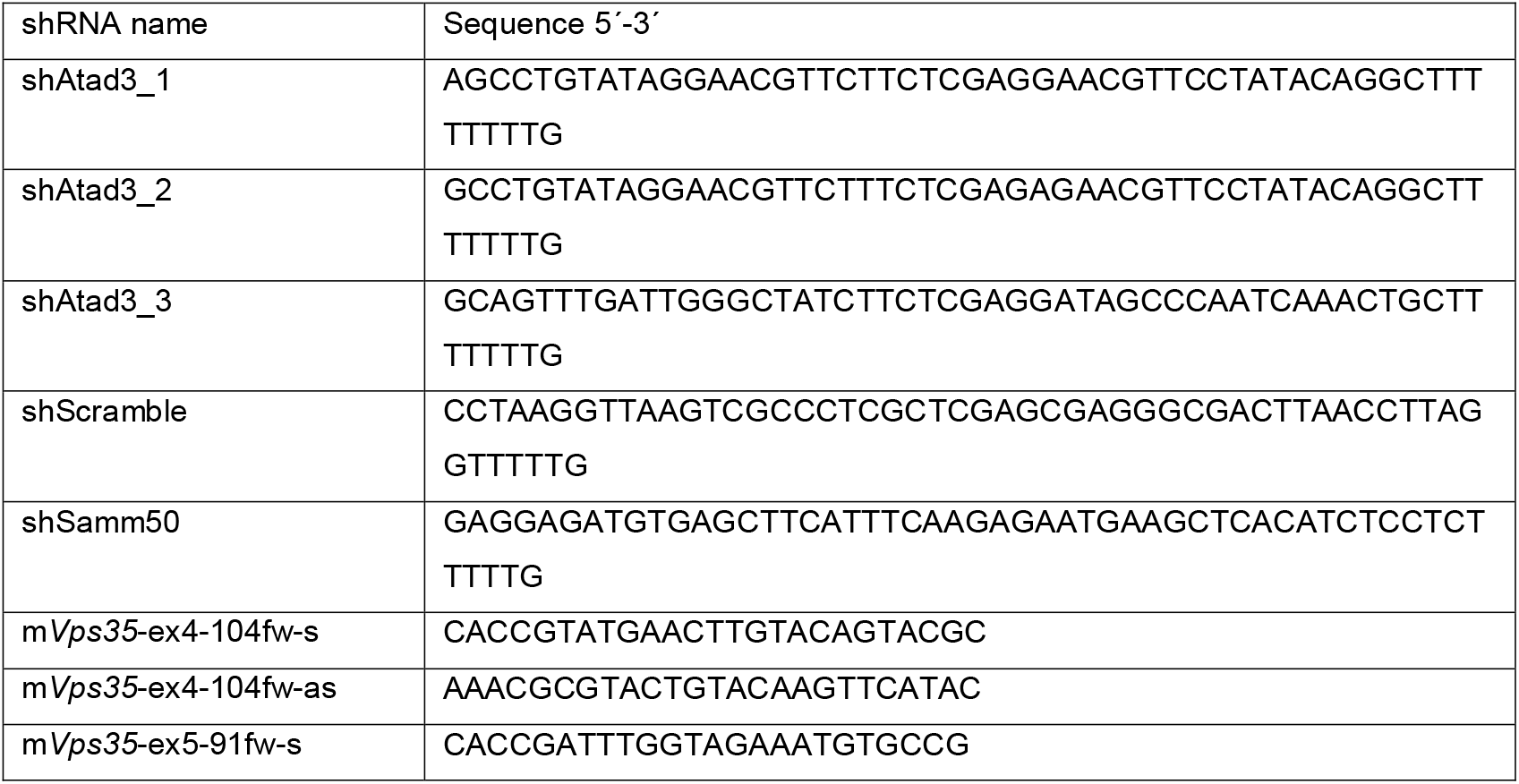
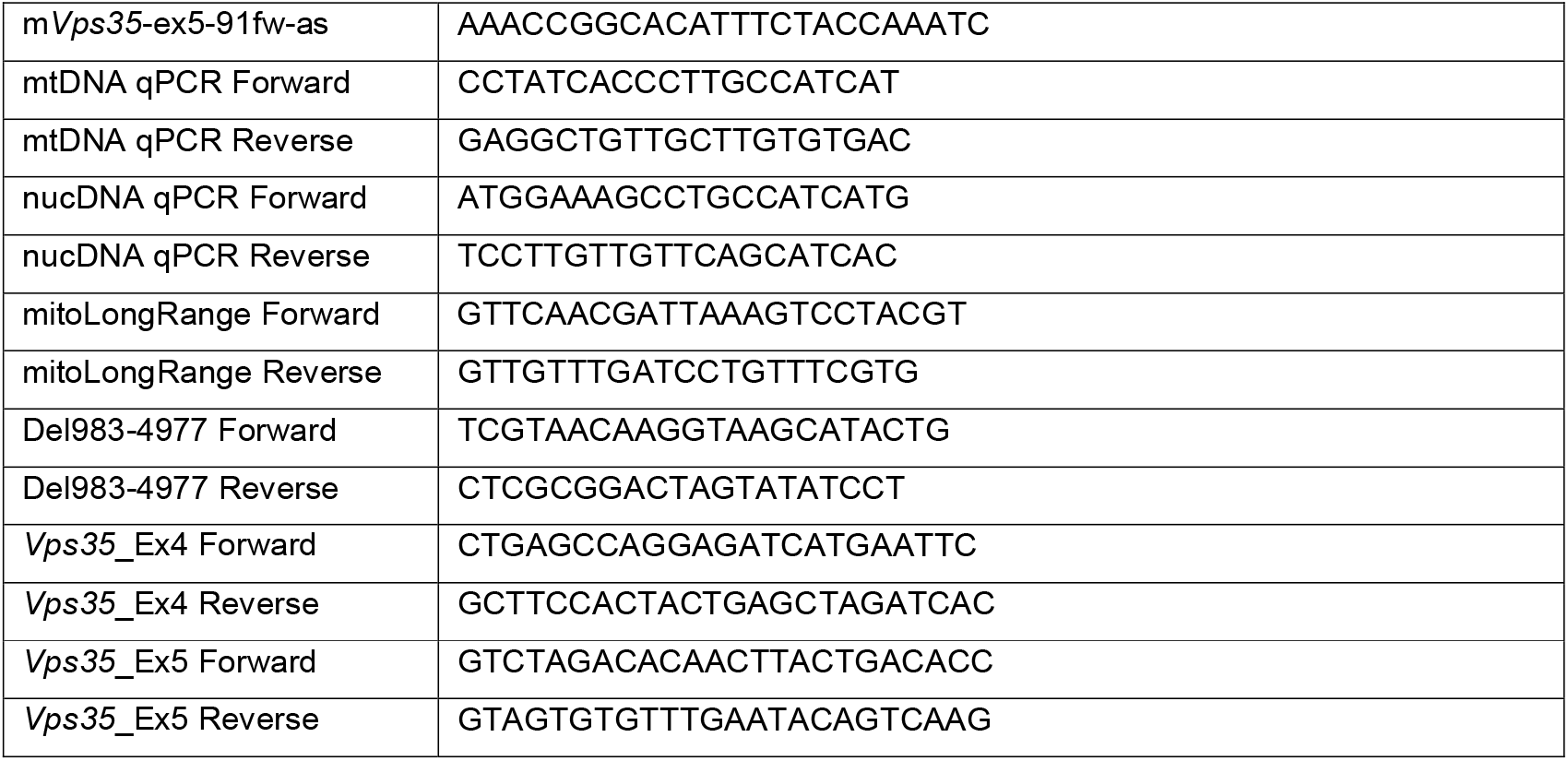
Oligonucleotides used in this study.

## Acknowledgements

We are thankful to CECAD Imaging and proteomics facilities for excellent technical support. We thank Nadine Niehoff and Katrin Lanz for technical assistance. We are grateful to Thomas Paß for critical review and discussion of the results. This work was supported by grants from the Deutsche Forschungs-gemeinschaft (PL 895/1-1) and Köln Fortune (341/2019) to DPM and RJW.

## Author contributions

Funding Acquisition, DPM and RJW; Conceptualization, DPM and RJW; Investigation and Formal Analysis, DPM, AS, SK, KM, JH; Resources JN; Analysis of MS Data, SK; Visualization, DPM; Writing, DPM and RJW; Writing-Review & Edit; DPM, RJW, SK, MK; Supervision, DPM and MK.

---

**Figure S1.**
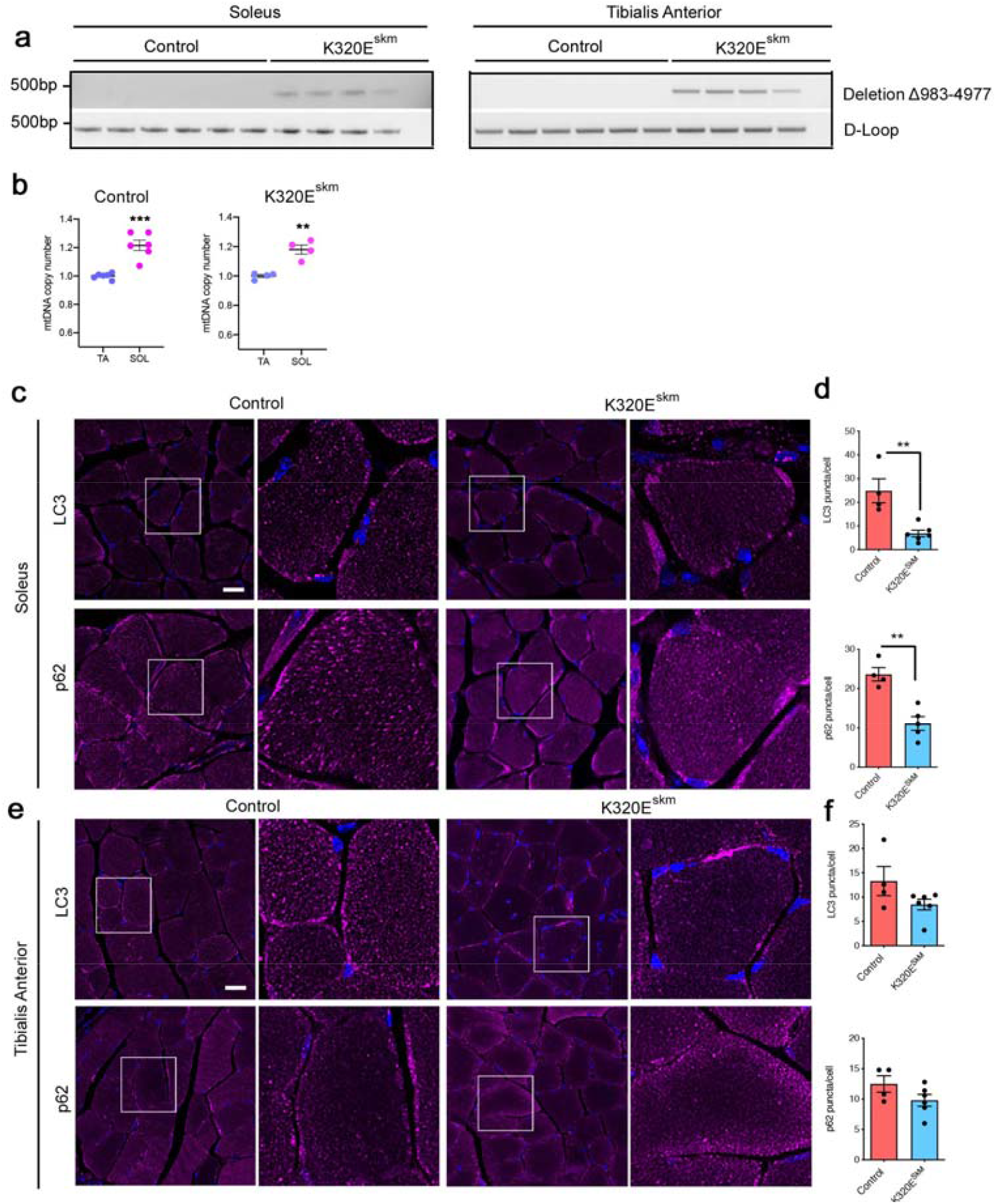
(**a**) Conventional PCR with specific oligonucleotides flanking the deletion mtDNA-Δ983-4977. (**b**) mtDNA copy number in muscles from 24 months old control and Twinkle-K320E mice. This graph shows a comparison between samples analysed in Fig 1b. (**c-f**) *In situ* immunofluorescence and image quantification showing autophagic markers LC3 and p62 in cryosections of M. soleus (**c, d**) and M. tibialis anterior (**e, f**). 5 random pictures with 4 fibers per picture were analysed per animal to obtain averaged values. Control: n=4; Twinkle-K320E: n=5. Bar, 20µm. (**b, c, e, h** and **i**) Unpaired Students’
s T-test. Mean ± SEM.

**Figure S2.**
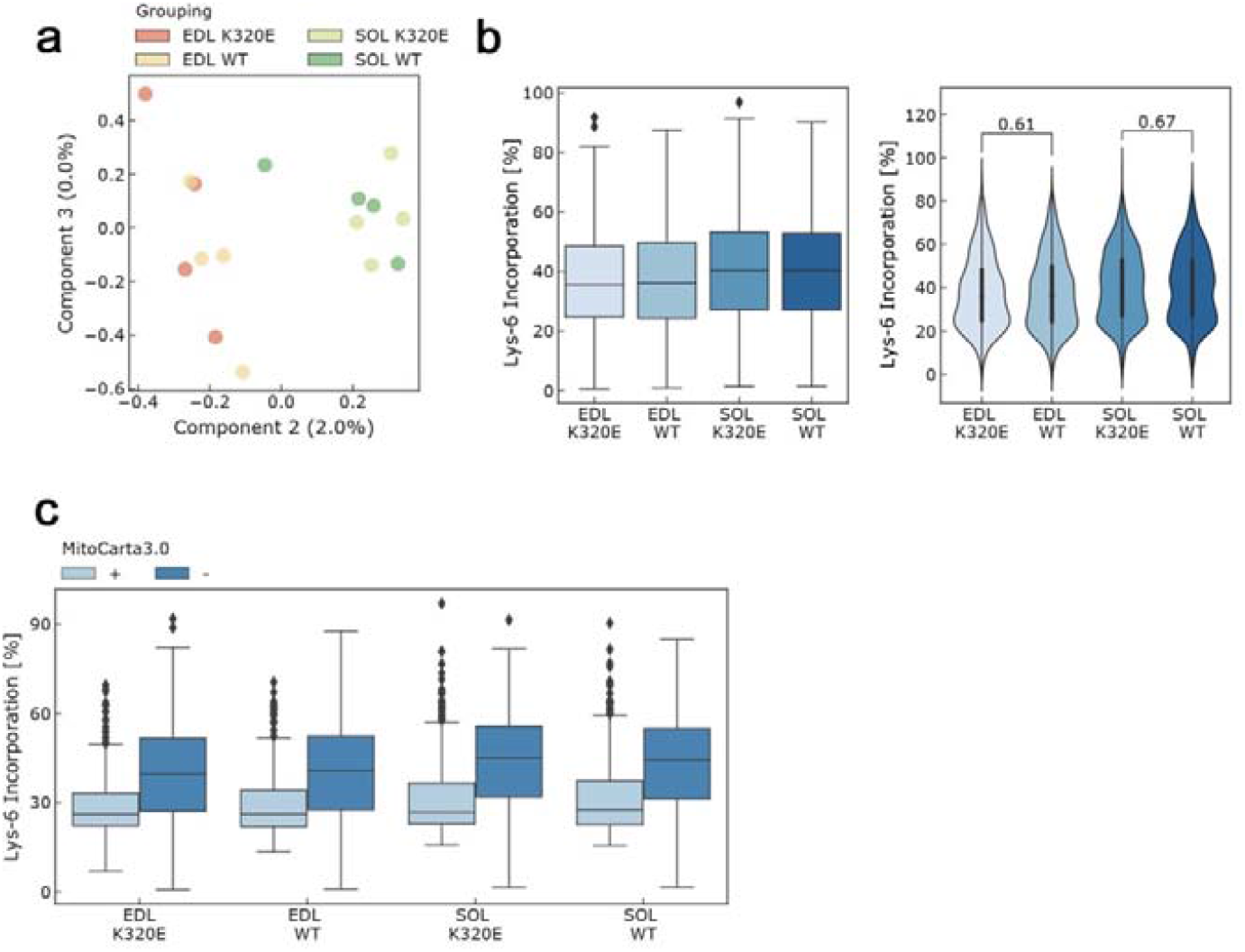
(**a**) Component analysis of *In vivo* Pulse SILAC in M. extensor digitorum longus (EDL) and M. soleus. (**b**) Box plot and Violin plots showing Lys-6 incorporation in muscles from Twinkle mice. (**c**) Box plot analysis for mitochondrial and non-mitochondrial proteins detected in the muscles analysed for *in vivo* Pulse SILAC. n=5.

**Figure S3.**
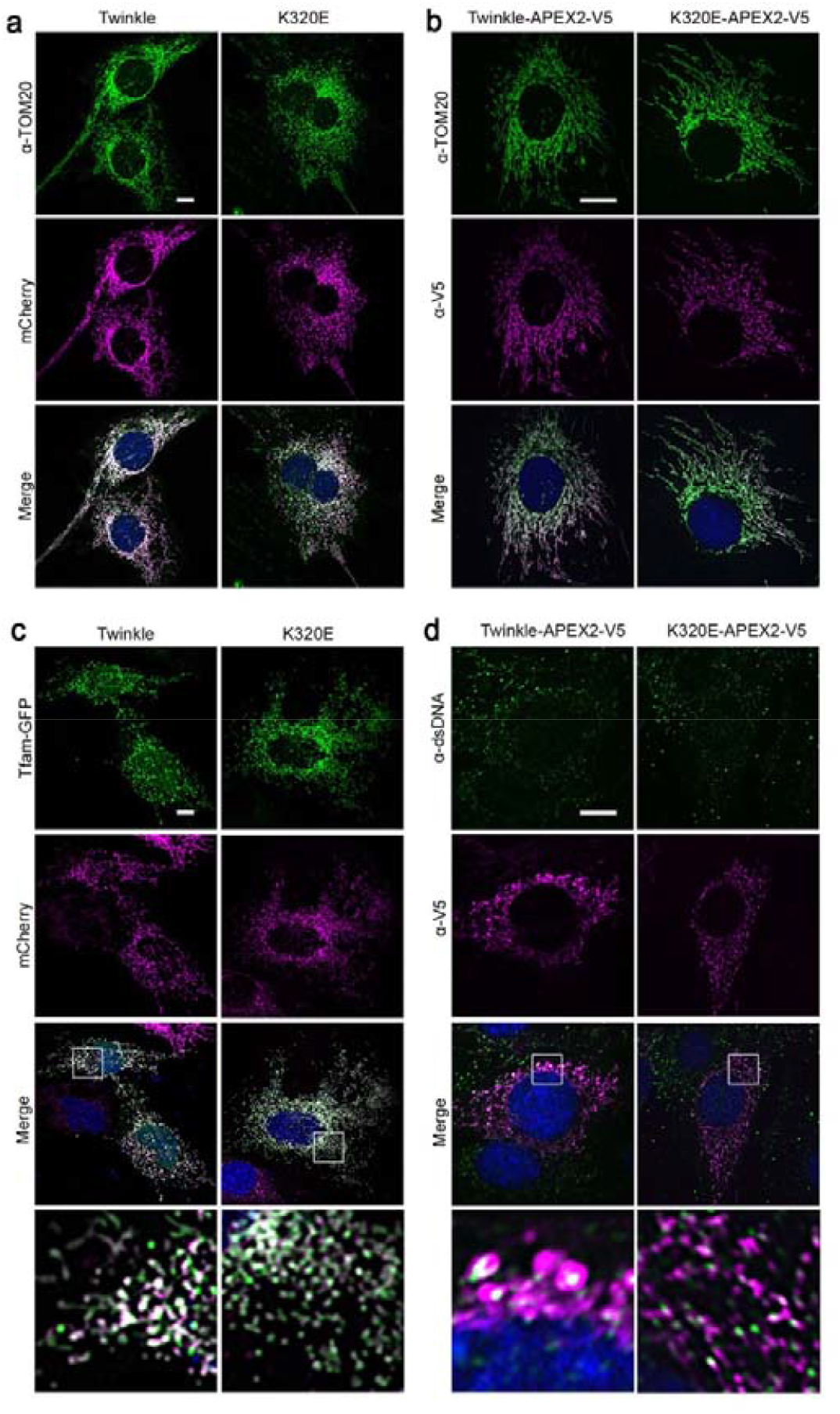
(**a, b**) α-TOM20 immunofluorescence of clones expressing (**a**) m-Cherry and (**b**) APEX2-V5 tagged Twinkle variants. (**c, d**) Immunofluorescence of (**c**) m-Cherry tagged clones transiently expressing Tfam-GFP and (d) APEX-V5 tagged clones probed with α-dsDNA antibody showing nucleoid localization (see magnifications in lower panels). Bar, 10 µm.

**Figure S4.**
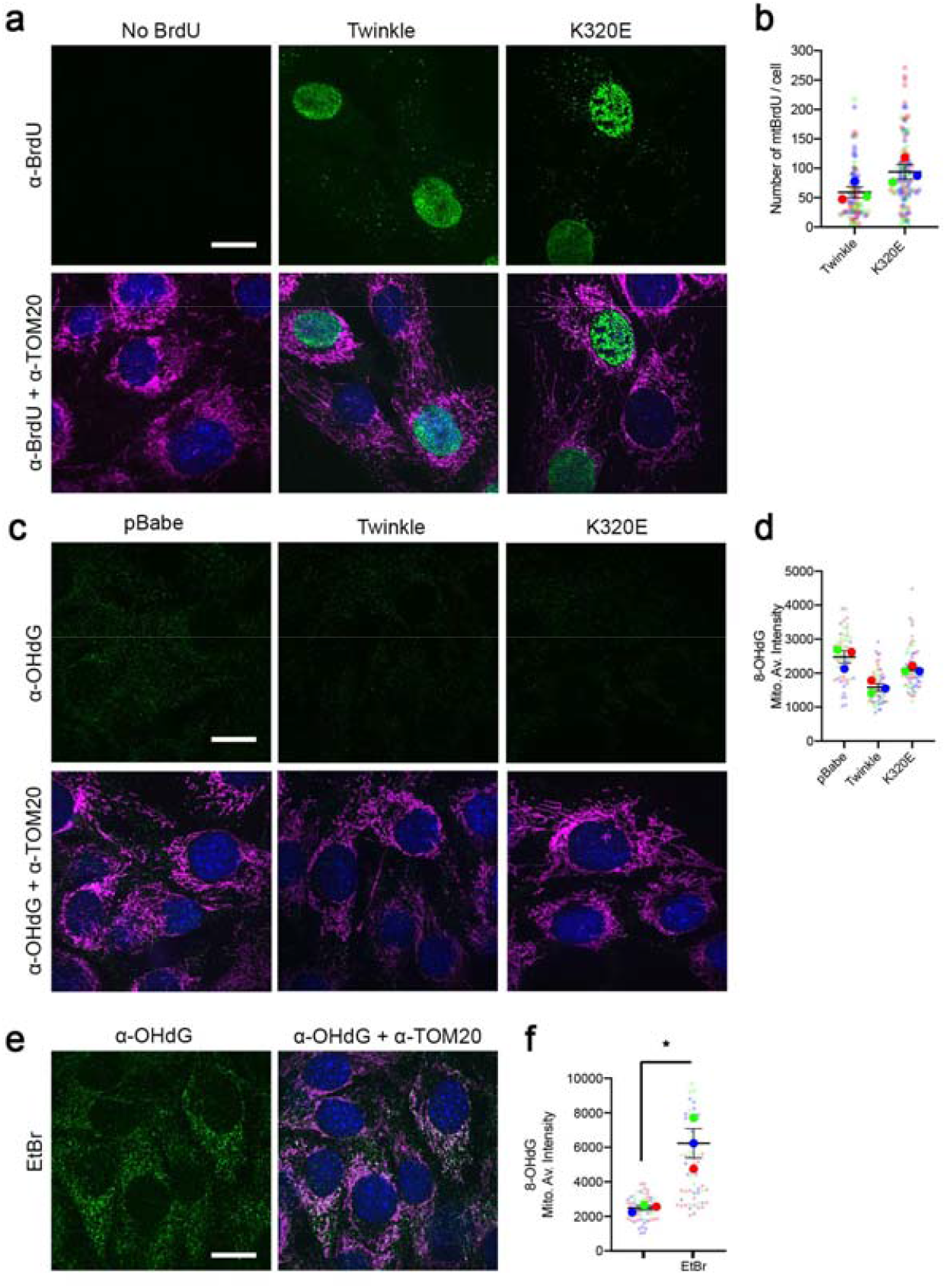
(**a, b**) Analysis of mtDNA replicating foci by α-BrdU and α-TOM20 immunofluorescence of C2C12 cells expressing Twinkle plasmids treated with 20 µM BrdU. Only BrdU foci inside the mitochondrial network were analysed. (**c, d**) α-OHdG and α-TOM20 immunofluorescence and quantification of the average intensity inside the mitochondrial network of C2C12 in steady state. (**e, f**) α-OHdG and α-TOM20 immunofluorescence and quantification of the average intensity inside the mitochondrial network in C2C12 treated 24h with 50 ng/ul EtBr. Students’
s T-test. Mean ± SEM. *, p<0.05. Scale Bar, 10 µm.

**Figure S5.**
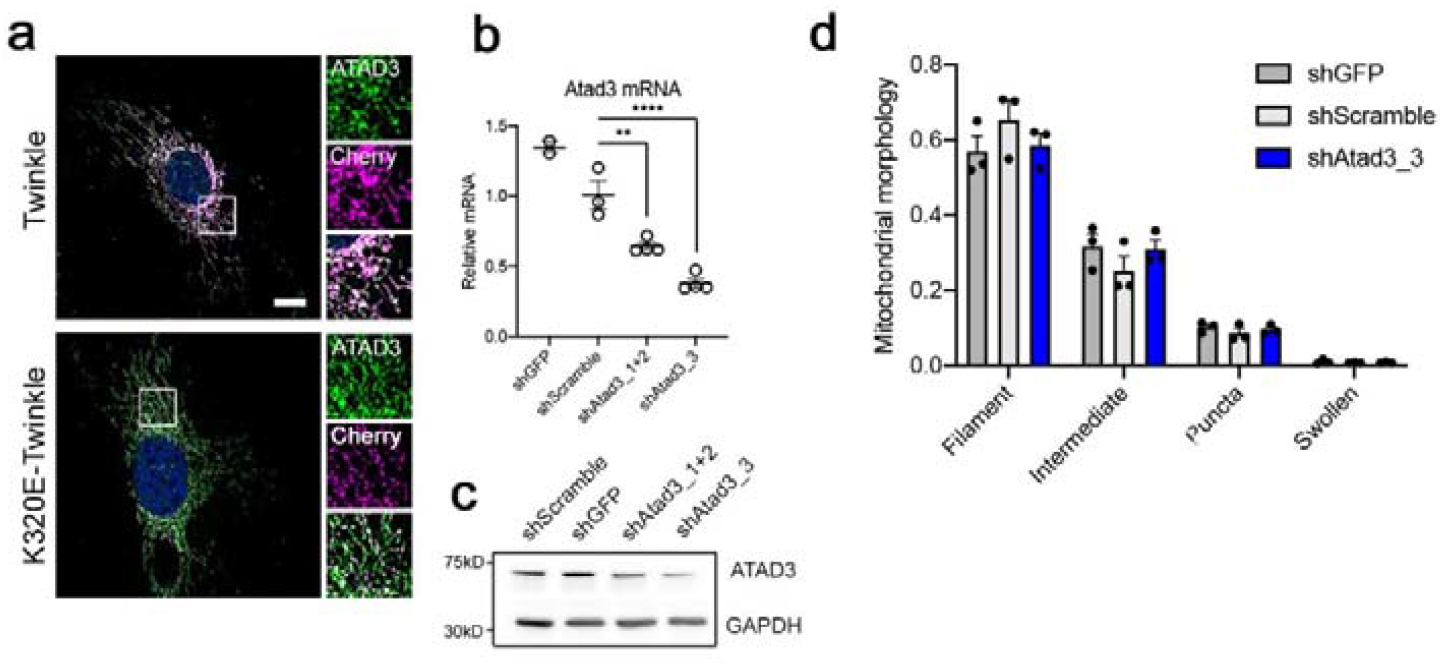
(**a**) Immunofluorescence of C2C12 cells expressing Twinkle-mCherry stained with α-ATAD3 antibody. Scale bar, 10 µm. (**b**) Analysis of Atad3 expression by RT-qPCR and (**c**) western blot upon shRNA transduction. n=3-4 (**d**) Quantification of Mitochondrial morphology in shATAD3 cells. n=3, >30 cells per replicate. (**b** and **d**) ANOVA, Tukey multiple comparison. **, p<0.01; ****, p<0.001. Mean ± SEM.

**Figure S6.**
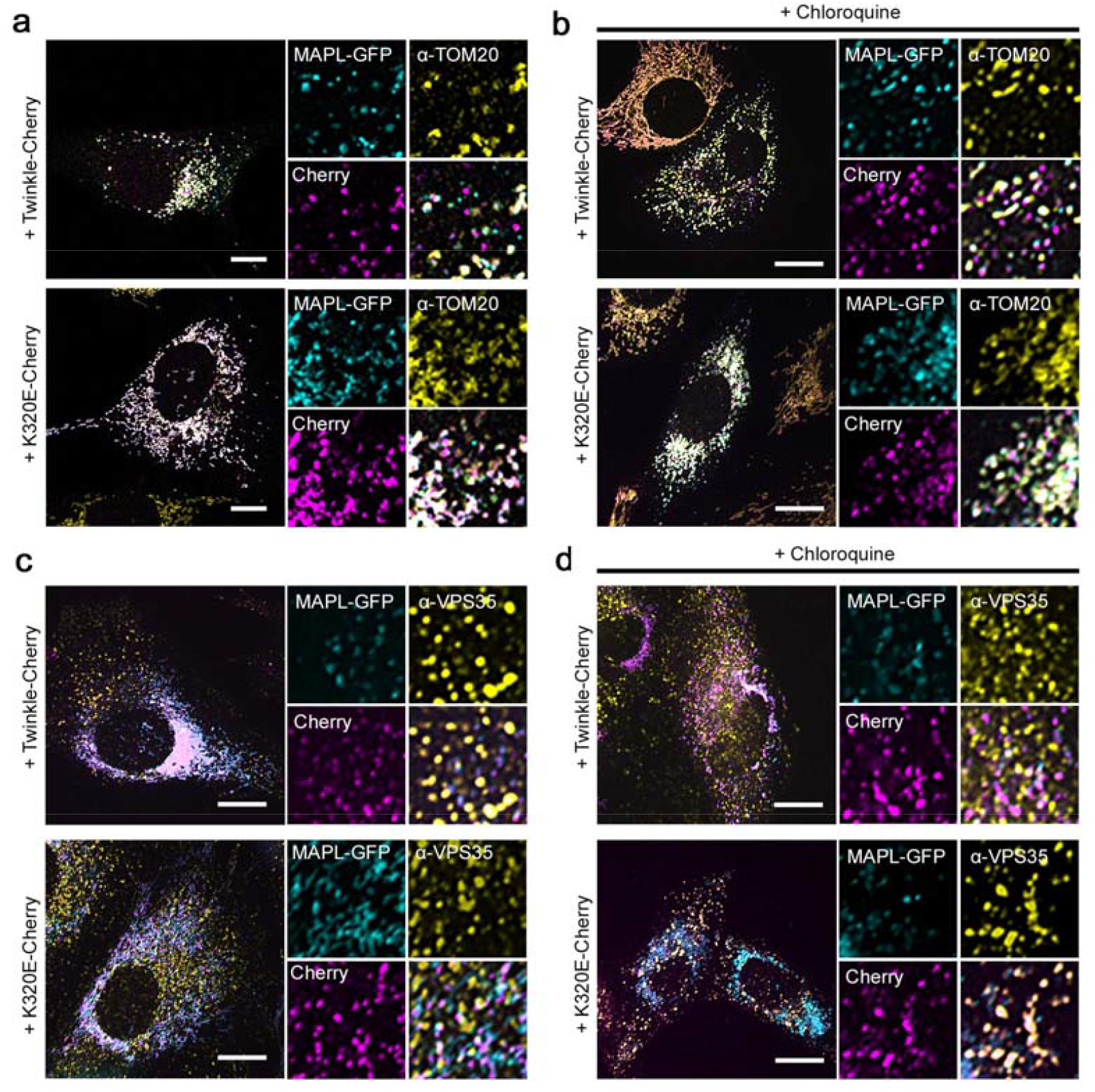
(**a-c**) Immunofluorescence of C2C12 cells expressing Twinkle-mCherry transiently transfected with MAPL-GFP and stained with α-TOM20 antibody (**a, b**) or α-VPS35 antibody (**c, d**). In (**b, d**) Cells were treated 4h with 10µM Chloroquine. Bar, 10 µm.

**Figure S7.**
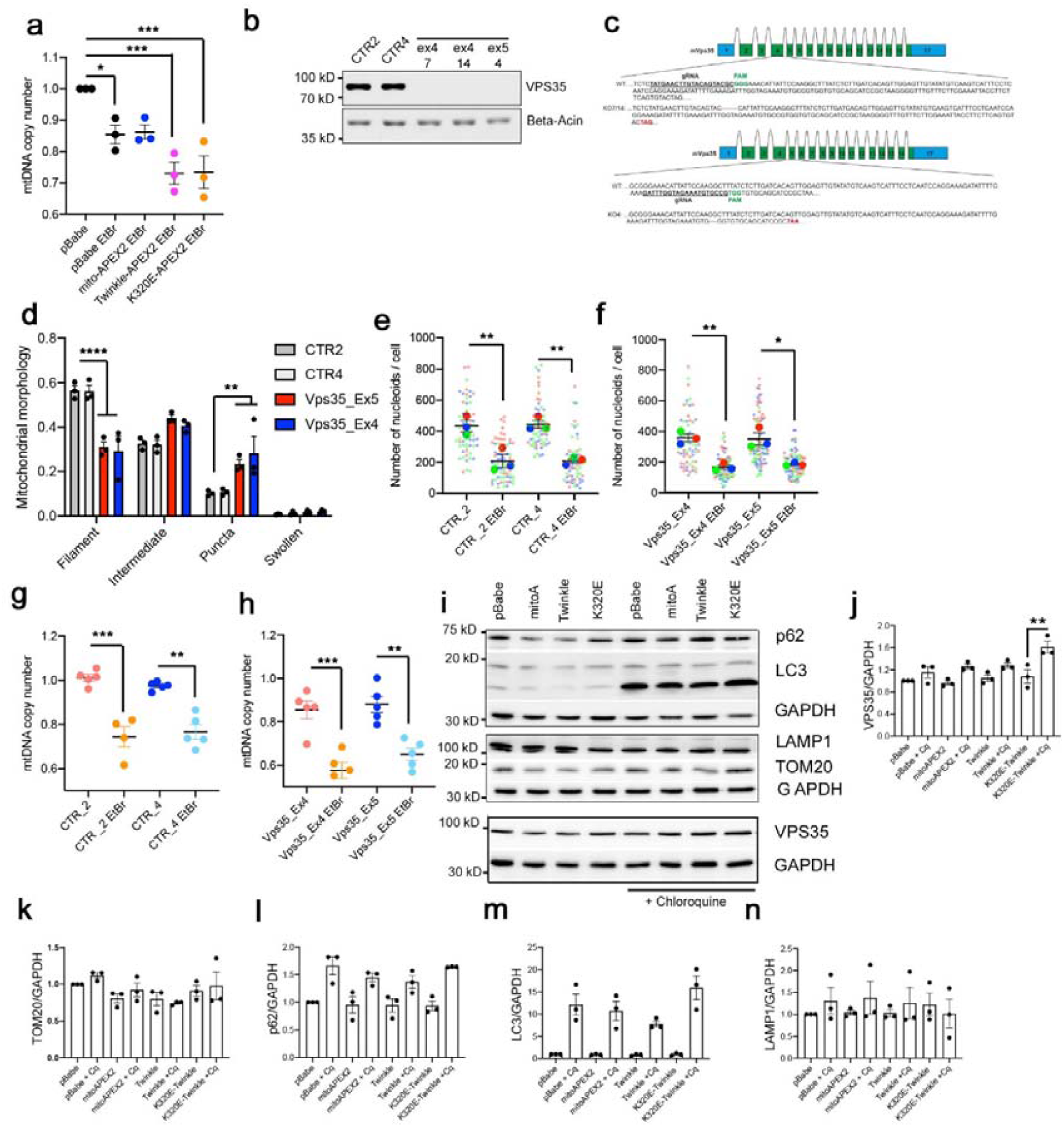
(**a**) mtDNA copy number in C2C12 cells treated for 7 days with 50ng/ml EtBr. n=3. (**b**) Western blot analysis of MEFs after CRISPR Cas9 mediated KO of *Vps35*. (**c**) Schematic representation of DNA modification in *Vps35* Exon 4 and Exon 5 triggered by CRISPR-Cas9. (**d**) Quantification of mitochondrial morphology in *Vps35* KO clones. n=3, >30 cells per replicate. (**e**) mtDNA foci quantification in control and (**f**) *Vps35* KO clones treated with EtBr. n=3. (**g**) qPCR mtDNA copy number analysis of control and (**h**) *Vps35* KO clones treated with EtBr. Students’
s T-test. **, p<0.01; ***, p<0.001. n=5. (**i**) Western blot analysis of autophagy markers and VPS35 levels in C2C12 cells expressing Twinkle-APEX2-V5 variants (WT, KE) or matrix targeted APEX2 protein, treated 24h with 10µM Chloroquine. n=3. (**j-n**) Intensity quantification of the indicated proteins relative to empty vector pB (pBABE). (**a, g, h and j-n**) ANOVA analysis of variance. Tukey multiple comparison test. Mean ± SEM. *, p<0.05; **, p<0.01; ***, p<0.001. (**d**) Two-way ANOVA. Control lines were compared with KO lines. **, p<0.01; ****, p<0.0001.

**Figure S8.**
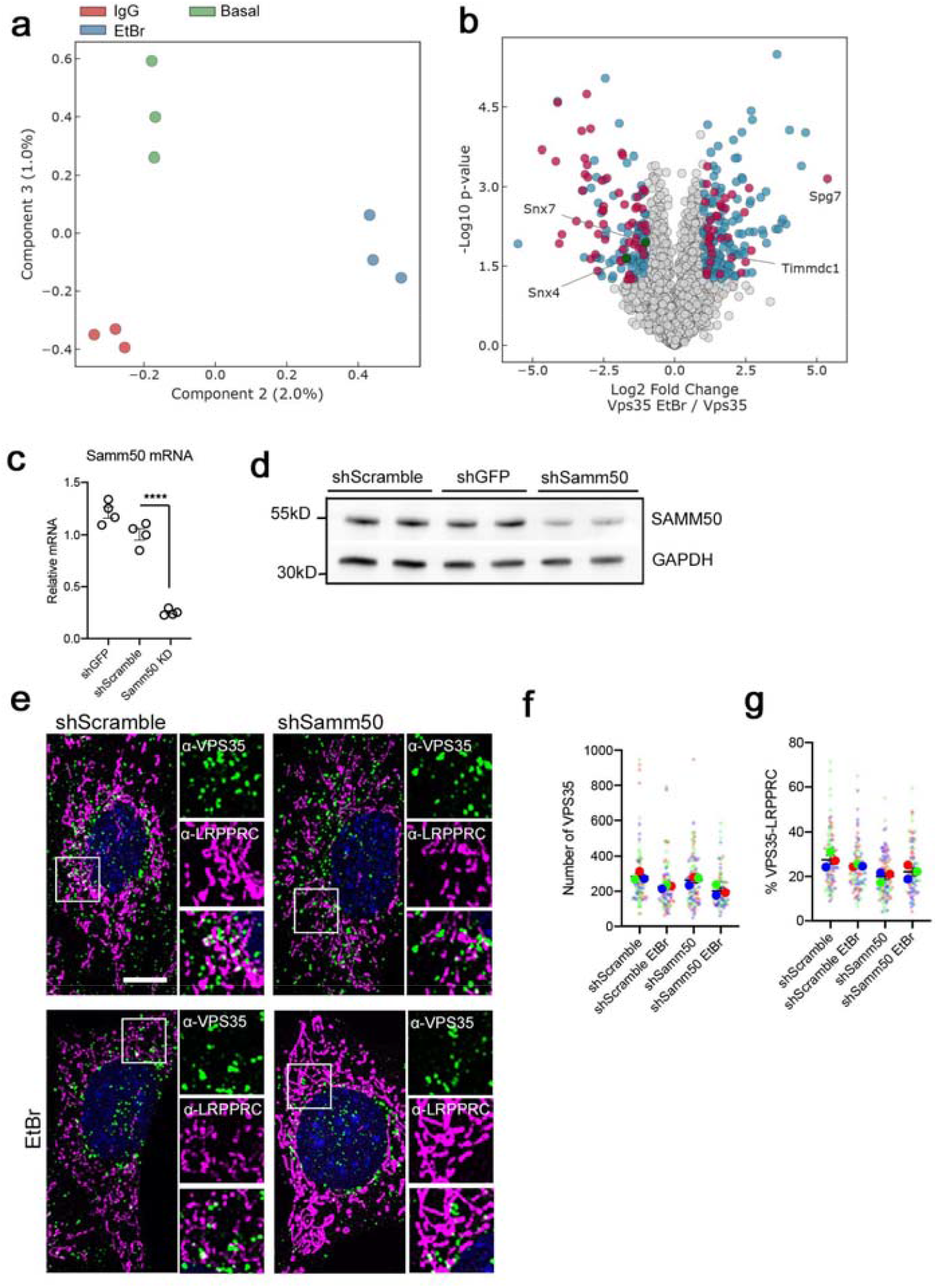
(**a**) Component analysis of VPS35 IP. (**b**) Comparison of VPS35 IP profiles from cells in basal medium or treated with 50ng/ml EtBr for 1 week. Only significantly different proteins are highlighted. Blue, cytosolic proteins; Red, mitochondrial proteins; Green, retromer proteins. (c) mRNA quantification and (d) western blot analysis of *Samm50* upon shRNA transduction. *Gapdh* mRNA and GAPDH was used as an internal control for both experiments. (**e**) Immunofluorescence of Control and Samm50 KD MEFs labelled with α-VPS35 and α-LRPPRC in basal and 7 days treated with 50ng/ml EtBr. Scale bar, 10µm. (**f, g**) Quantification of VPS35 particles and VPS35 in contact with LRPPRC. n=3. Mean ± SEM. ANOVA analysis of variance. Tukey multiple comparison test.

**Figure S9.**
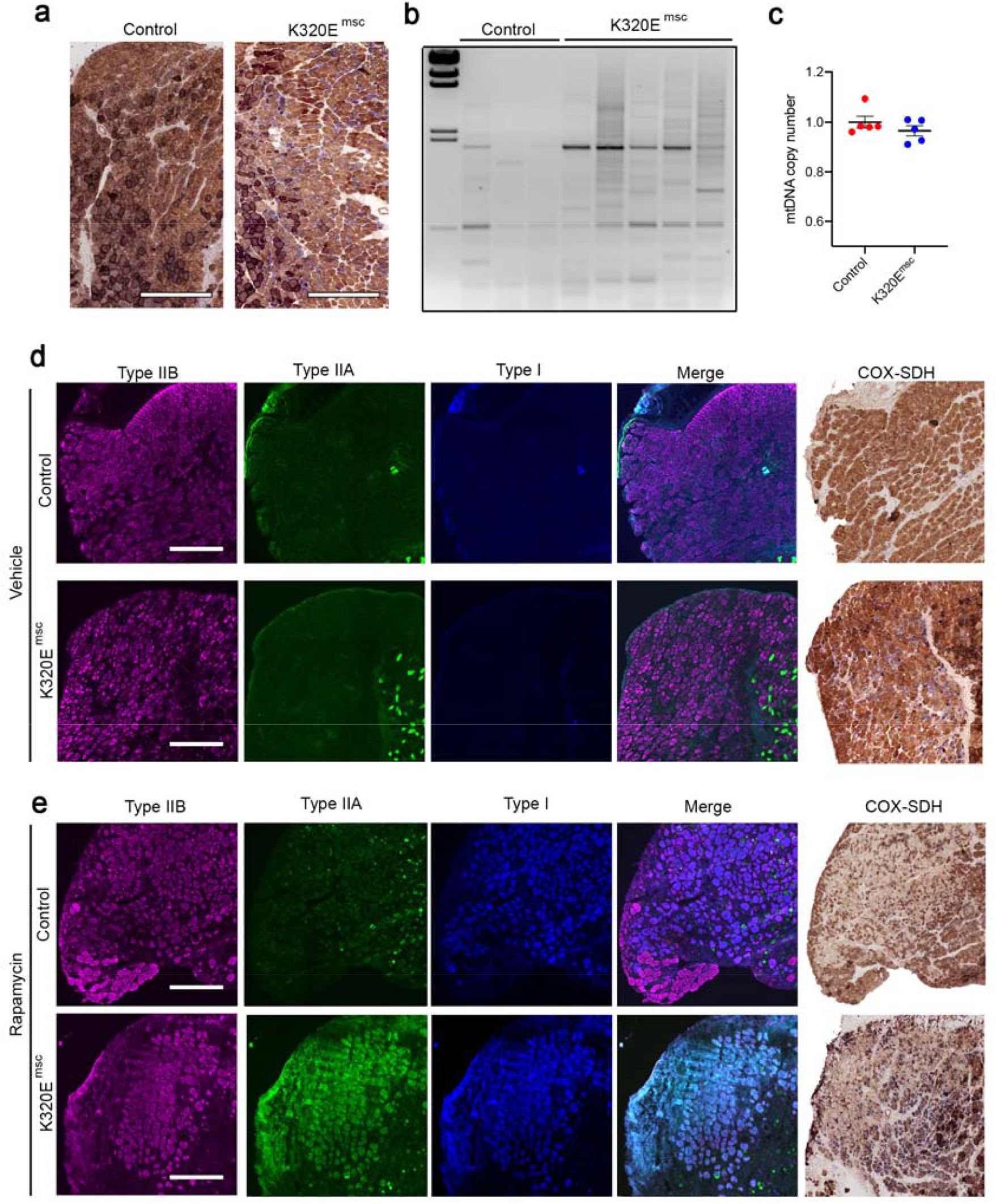
(**a**) Representative image for COX-SDH staining in a regenerated area from the M. Tibialis anterior from control and K320E-Twinkle^msc^ -mice. Bar, 500 µm. (**b**) Long Range PCR for mtDNA deletions and, (**c**) mtDNA copy number analysis in control and Twinkle-K320E mice. n=5. Muscle regeneration was induced with intramuscular injection of 10µM Cardiotoxin (*Naja Pallida*). Students’
s T-test. (**d**) Fiber type and COX-SDH serial analysis of regenerated muscles in vehicle and (**e**) rapamycin treated mice. Mean ± SEM. Bar, 250 µm.

